# Symbiosis genes show a unique pattern of introgression and selection within a *Rhizobium leguminosarum* species complex

**DOI:** 10.1101/526707

**Authors:** Maria Izabel A. Cavassim, Sara Moeskjær, Camous Moslemi, Bryden Fields, Asger Bachmann, Bjarni J. Vilhjalmsson, Mikkel H. Schierup, J. Peter W. Young, Stig. U. Andersen

**Author notes:** Authors for correspondence: Stig Uggerhøj Andersen, J. Peter W. Young.

## Abstract

Rhizobia supply legumes with fixed nitrogen using a set of symbiosis genes. These can cross rhizobium species boundaries, but it is unclear how many other genes show similar mobility. Here, we investigate inter-species introgression using *de novo* assembly of 196 *Rhizobium leguminosarum* bv. *trifolii* genomes. The 196 strains constituted a five-species complex, and we calculated introgression scores based on gene tree traversal to identify 171 genes that frequently cross species boundaries. Rather than relying on the gene order of a single reference strain, we clustered the introgressing genes into four blocks based on population structure-corrected linkage disequilibrium patterns. The two largest blocks comprised 125 genes and included the symbiosis genes, a smaller block contained 43 mainly chromosomal genes, and the last block consisted of three genes with variable genomic location. All introgression events were likely mediated by conjugation, but only the genes in the symbiosis linkage blocks displayed overrepresentation of distinct, high-frequency haplotypes. The three genes in the last block were core genes essential for symbiosis that had, in some cases, been mobilized on symbiosis plasmids. Inter-species introgression is thus not limited to symbiosis genes and plasmids, but other cases are infrequent and show distinct selection signatures.

## Introduction

Mutation and meiotic recombination are the main sources of genetic variation in eukaryotes. In contrast, prokaryotes can rapidly diverge through other types of genetic exchange collectively known as horizontal gene transfer (HGT). These include transformation (through the cell membrane), transduction (through a vector), and conjugation (through cell-to-cell contact) [Ochman and Lawrence 2000; Hanage 2016]. These processes can introgress adaptive genes to distantly related species, creating specific regions of high genetic similarity.

It has often been suggested that HGT would blur the boundaries between species to the extent that species phylogenies would be better represented by a net-like pattern than a tree [Doolittle 1999]. This notion may have arisen when the methods for studying prokaryotic evolution and species delineation were still rudimentary, making it challenging to accurately evaluate the rate of HGT [Konstantinidis and Tiedje, 2007]. With ever-increasing numbers of bacterial whole-genome sequences (WGS), it has become possible to re-evaluate bacterial species classification [Konstantinidis et al., 2006; Jain et al., 2018]. HGT, or introgression, events can be inferred using parametric or phylogenetic methods. Parametric methods rely on comparing gene features such as nucleotide composition or *k*-mer frequencies to a genomic average in order to detect outliers, which may be associated with HGT [Daubin et al., 2003; Burge and Karlin 1995]. Explicit phylogenetic methods are based on comparisons of gene and species trees aimed at detecting topological differences, whereas implicit phylogenetic methods rely on detecting aberrant distances to an outgroup reference [Lerat et al., 2003].

Whether sympatric species frequently exchange genetic material through HGT is still an open question and the nitrogen-fixing symbiont of legumes, *Rhizobium leguminosarum*, is a useful model for investigating inter-species introgression through HGT. There is extensive literature documenting the sharing of symbiosis-related genes among distinct, and sometimes distant, species of rhizobia [Rogel et al., 2011; Remigi et al., 2016; Andrews et al., 2018]. This occurs whether the genes are on plasmids [Segovia et al., 1993; Haukka et al., 1998; Laguerre et al., 2001; Pérez-Carrascal et al., 2016] or on conjugative chromosomal islands [Sullivan et al., 1995; Nandasena et al., 2006]. It has been previously observed that *R. leguminosarum* can be divided into distinct genospecies (gsA, gsB, gsC, gsD and gsE), but the host-specific symbiovars that nodulate white clover (*R. leguminosarum* sv. *trifolii*) and vetch (*R. leguminosarum* sv. *viciae*) are not confined to distinct genospecies [Kumar et al. 2015]. This provides another example of symbiosis gene transfer between sympatric rhizobia.

Symbiosis genes are known to increase the fitness of both symbiont and plant host (reviewed by Friesen 2012), but it is still unclear if their frequent introgression represents a special case, or if HGT is common for a wide range of genes in sympatric rhizobia. To address this question and obtain a more general understanding of introgression characteristics and mechanisms among sibling bacterial species, we assembled 196 *R. leguminosarum* genome sequences and carried out an unbiased introgression analysis.

## New approaches

### Detection of gene introgression

We introduce a simple and robust method to detect genes that show introgression across species boundaries. We build high-quality groups of orthologous genes, align them and then traverse each resulting gene tree, counting how many times an interspecies transition is encountered. This provides a direct measure of introgression frequency.

### Intergenic Linkage Disequilibrium (LD) analysis corrected for population structure

In order to identify blocks of linked genes against a background of variable gene order in a bacterial species complex, we develop a method for quantifying intergenic LD while correcting for population structure. We first calculate the genetic relationship matrix of all SNPs and use it to generate pseudo-SNPs corrected for population structure (Mangin et al., 2012). We then calculate intergenic LD by applying Mantel tests to pseudo-SNP genetic relationship matrices for pairs of genes.

## Results

### Five distinct species constitute a *R. leguminosarum* species complex

Previous work has shown the existence of five distinct *R. leguminosarum* genospecies within one square meter of soil [Kumar et al., 2015]. To acquire a broader diversity sample from a wider geographical area, we isolated 196 rhizobium strains from white clover root nodules harvested in Denmark, France and the UK (**Figure S1-2, Table S1**). We then sequenced and *de novo* assembled the genomes of all 196 strains, followed by genome annotation and construction of orthologous gene groups (**Figure S3-5, Table S2**). To determine the relationship of our 196 strains with the previously identified genospecies, we constructed a phylogenetic tree containing RpoB sequences from known representatives of the five genospecies in addition to those from our 196 strains. This allowed us to assign all of our strains to a specific genospecies based on their position in the tree (**Figure S6**). Since our extended sampling did not result in identification of additional genospecies, the five genospecies, gsA-E, likely represent a large part of northern European *R. leguminosarum* diversity.

The 196 strains shared a total of 4,204 core gene groups, which had a higher median GC content than the 17,911 accessory gene groups (**Figure S7**). We calculated average nucleotide identity (ANI) based on 305 conserved genes (**Supplementary Table S3**) and on 6,529 genes present in at least 100 strains and clustered the strains based on pairwise ANI (**Figure 1a-b**). The strains were collected from different countries and field management regimes, but they clustered mainly by genospecies, although substructure related to geographic origin was also evident (**Figure 1a-b**). These patterns were similar when clustering was carried out based on shared gene content (**Figure 1**, **Figure S7**). In conjunction with earlier evidence that the standard whole-genome measure of ANI was lower than 0.95 for inter-genospecies comparisons [Kumar et al. 2015], these results confirm that the genospecies should be considered genuinely distinct species constituting an *R. leguminosarum* species complex (**Figure 1a**).

**Figure 1.**
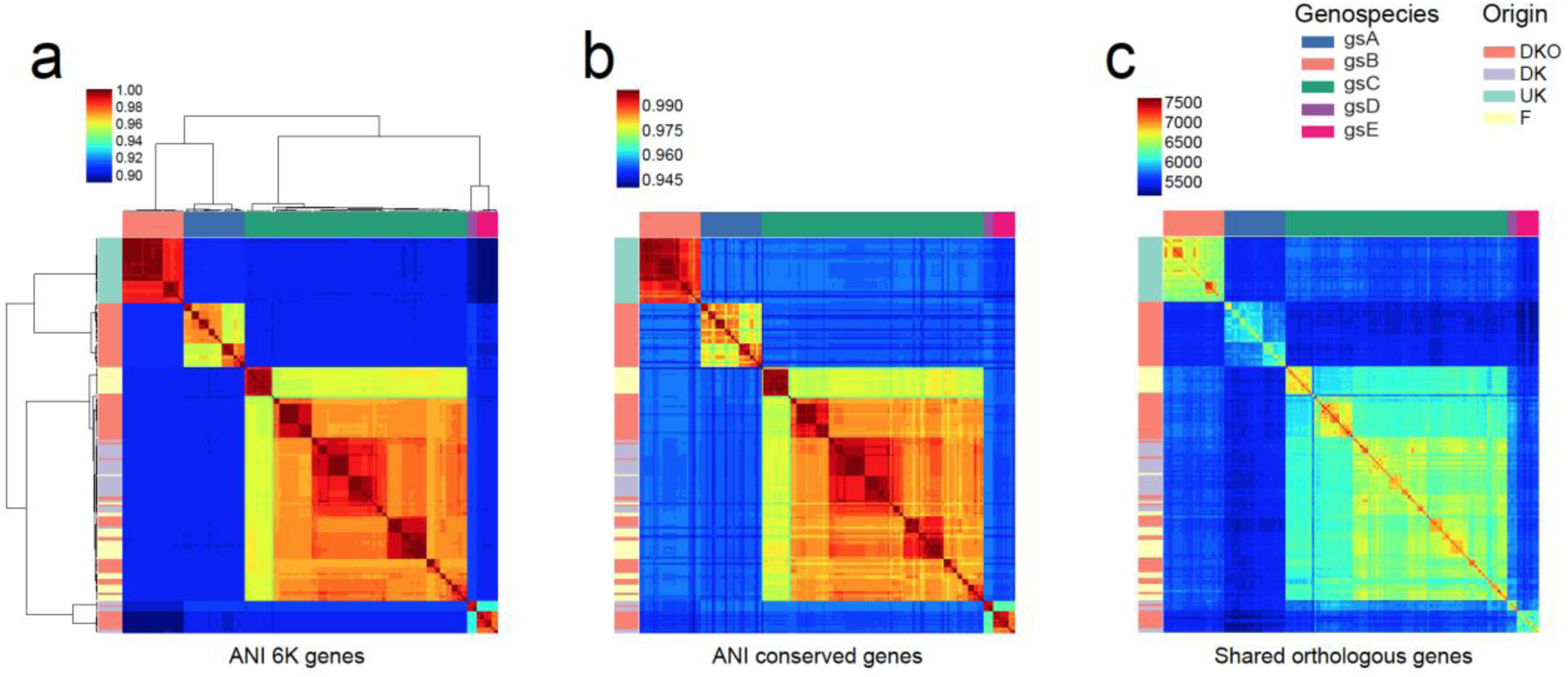
Genetic divergence across 196 rhizobium strains. Pairwise comparisons of genetic diversity were analyzed at different levels. (**a**) Proportion of shared single nucleotide polymorphisms (SNPs) in genes that were present in at least 100 strains and that passed filtering criteria (6,529 genes, 441,287 SNPs). Clusters of strains with SNP identity above 96% were recognised as 5 genospecies: gsA (blue), gsB (salmon), gsC (green), gsD (purple), gsE (pink) as indicated in the legend. **(b)** Average nucleotide identity for concatenated sequences of 305 housekeeping genes. **(c)** Number of shared genes. Strain origins are indicated by coloured bars at the left (DKO in red, DK in purple, F in yellow, and the UK in green). Strains were ordered by clustering of the SNP data.

### Plasmids are not genospecies specific

The genome of *R. leguminosarum* consists of a chromosome and a variable number of low-copy-number plasmids, including two that can be defined as chromids due to their size, ubiquitous presence across strains, and core gene content [Kumar et al. 2015; Young et al. 2006; Harrison et al., 2010]. In order to characterize the plasmid diversity within this species complex, we examined the sequence variation of a plasmid partitioning gene (*repA*) that is essential for stable maintenance of nearly all plasmids in *Rhizobium*. From all 196 genomes, 24 distinct *repA* sequence groups were identified. However, four of these correspond to isolated *repA*-like genes that are not part of *repABC* operons, and twelve others were rare (in no more than four genomes), so eight *repA* types account for nearly all plasmids identified (**Figure 2; Table S4**). We numbered them Rh01 to Rh08 in order of decreasing frequency in the set of genomes. Of these, Rh01 and Rh02 correspond to the two chromids *pRL12* and *pRL11* of the reference strain 3841 [Young et al., 2006] and are present in every genome. The distribution of the other plasmids shows some dependence on genospecies, but none are confined to a single genospecies. For example, Rh03 is present in all strains of gsA, gsB and gsC, but absent from gsE and in just one gsD strain, while Rh05 is universal in gsA and gsB but absent elsewhere (**Figure 2; Table S5**).

**Figure 2.**
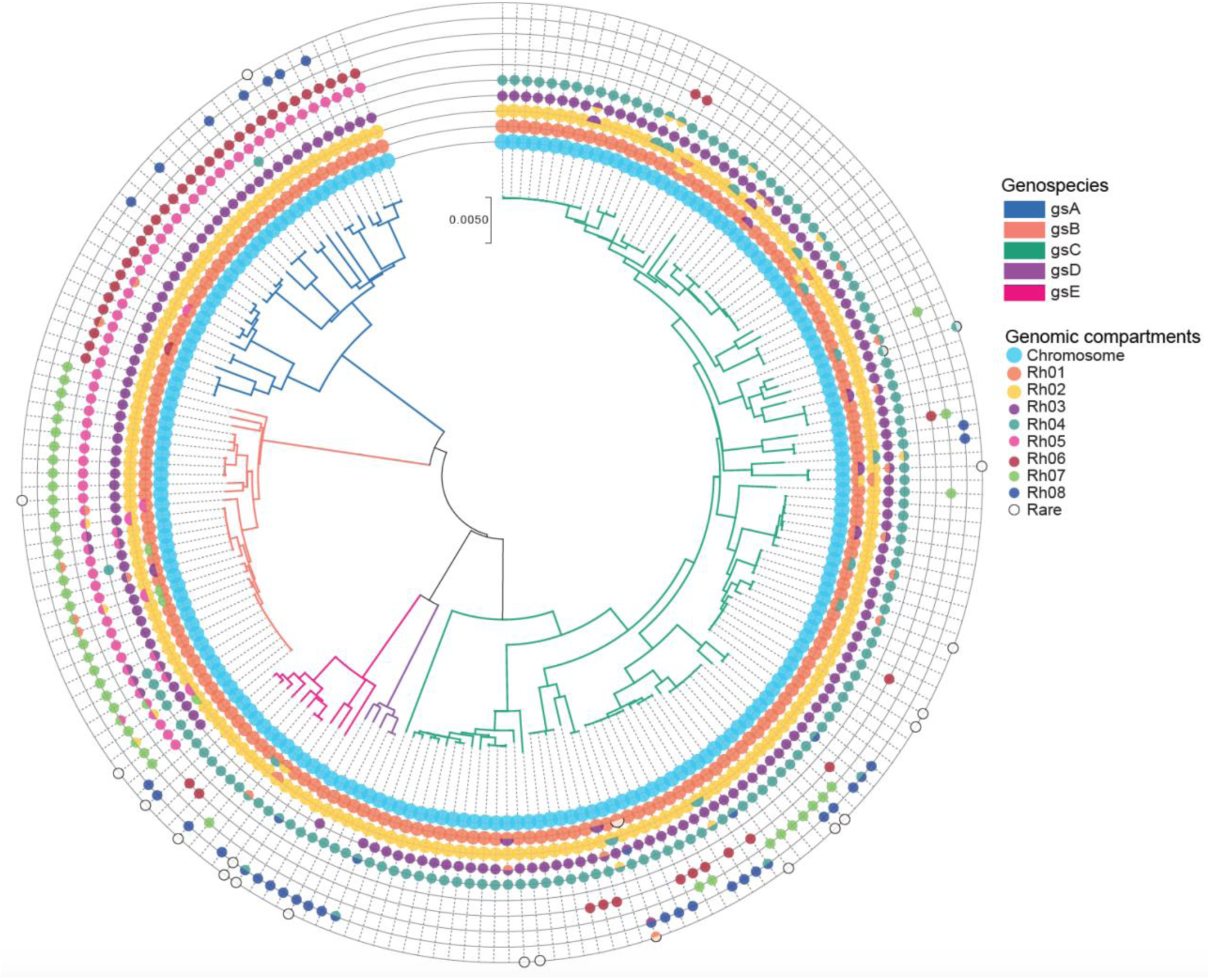
Characterization of plasmid diversity. Species phylogeny based on the concatenation of 305 core genes using the neighbor-joining method. Branches are coloured by genospecies. Circles represent the genomic compartments observed in each strain. Chromids (Rh01 and Rh02) and plasmids (Rh03, Rh04, Rh05, Rh06, Rh07, Rh08) were defined based on the genetic similarity of the RepA plasmid partitioning protein.

### Identification of introgression events based on gene trees

To evaluate the general rate of HGT within the present *R. leguminosarum* species complex, we developed a method to detect and quantify introgression (see Material and Methods). For a given gene, present in all five genospecies, traversal of the gene phylogeny should encounter only four inter-species transitions if no introgression had occurred, yielding an introgression score of 0. All transitions in addition to the four expected would indicate introgression events, adding to the introgression score (**Figure S8**). Most gene groups displayed very low introgression scores of 0 and 1 (**Figure 3a**), showing that introgression events were generally rare. At the other extreme of the distribution, we identified 171 genes with an introgression score above 10, indicating that they relatively frequently cross species boundaries (**Figure 3b**).

**Figure 3.**
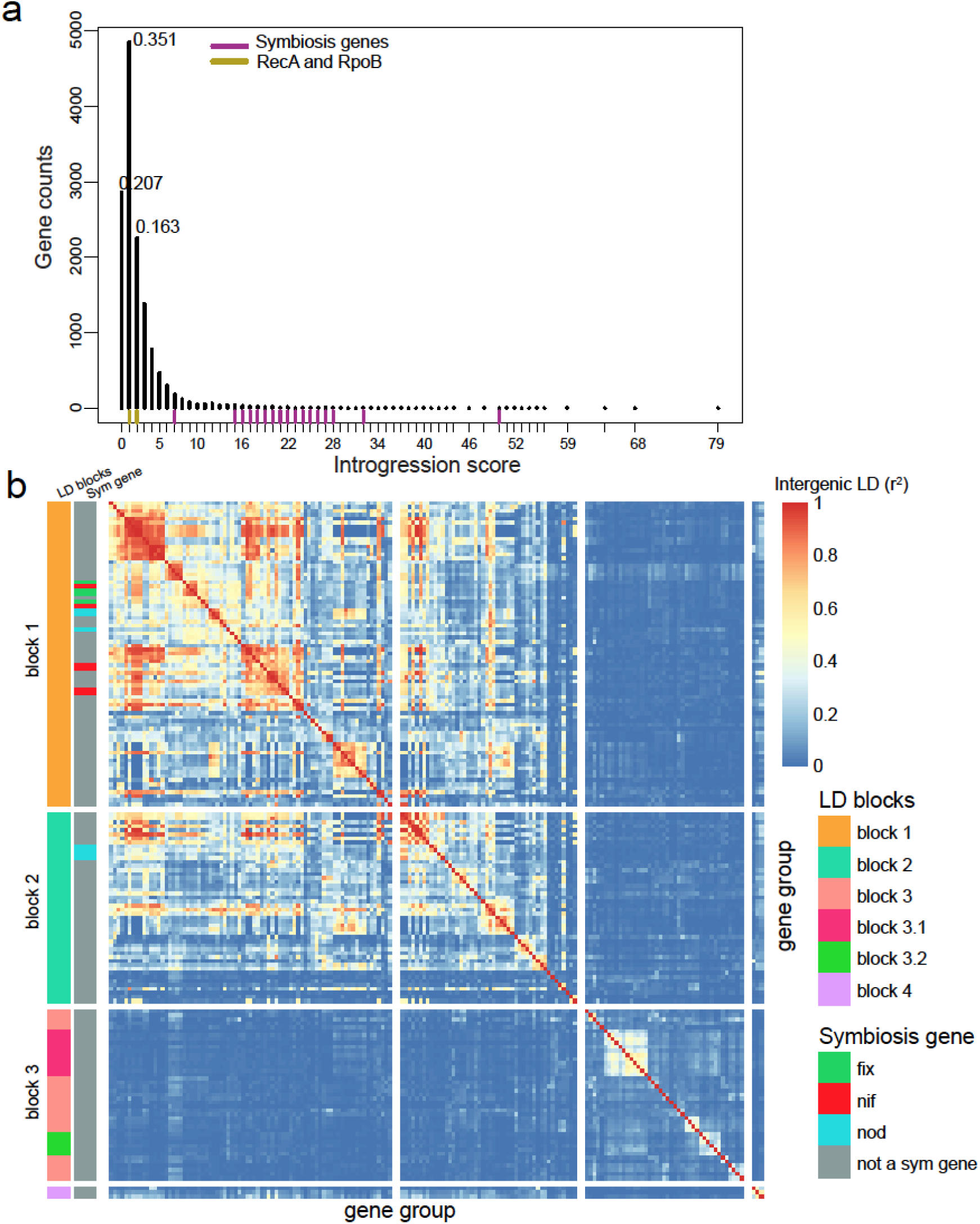
Introgression score values and its genomic distribution. **(a)** Distribution of introgression scores based on genes present in at least 2 genospecies (13,843)**. Pairwise Intergenic LD across highly introgressed genes (171 genes).** The x and y axes represent genes ordered by LD clustering rather than physical position. Genes were classified by LD blocks and by their contribution to symbiosis. The warmer the color the greater the intergenic correlation (r^2^). Chromosomal genes are found in LD block 3, while plasmid-borne genes are clustered in the first two blocks.

### Clustering genes using population structure-corrected LD

Gene order was variable across the species complex (**Figure S9**). To understand the nature of the genes displaying introgression, we therefore grouped them by linkage disequilibrium patterns rather than relying on the gene order of a single reference strain. The Mantel test is used to compare pairs of distance matrices, and here we used it to calculate intergenic LD by comparing genetic relationship matrices (GRM) [VanRaden, 2008]. However, when population structure exists, this approach suffers from inflation [Guillot and Rousset, 2013], and we observed this effect in our data as unexpectedly high levels of LD between plasmid-borne symbiosis genes and chromosomal core genes (**Figure S11a**). To address this issue, we calculated a genetic relationship matrix based on all SNPs and used it to generate pseudo-SNPs corrected for population structure for every gene (see Material and Methods, Mangin et al., 2012). We then compared the gene pseudo-SNP genetic relationship matrices using the Mantel test in order to calculate intergenic LD. After this correction for population structure, symbiosis genes and chromosomal core genes no longer appeared to be in LD (**Figure S11b**). We then proceeded to cluster the 171 genes that frequently crossed species boundaries based on their LD patterns. The genes separated into four clusters, where LD blocks 1 and 2 comprised the plasmid-borne symbiosis genes, block 3 contained mainly chromosomal genes and block 4 comprised three genes with a distinct LD pattern (**Figure 3b, Figure S10**). It is worth noting that, because of our stringent criteria, the LD blocks detected by our method are just a representation of some of the introgressed genes within each LD block region (**Figure 4a-c**).

**Figure 4.**
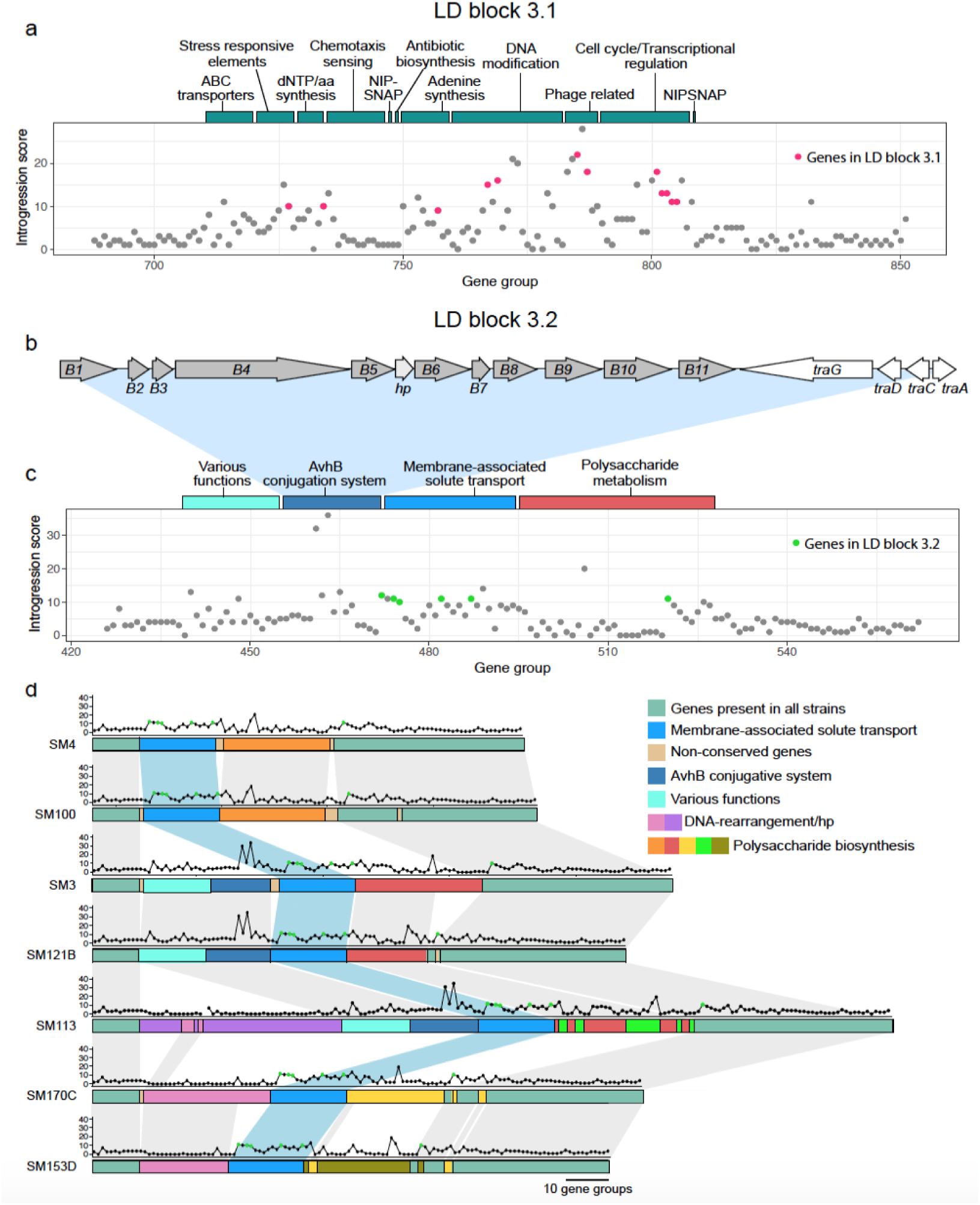
Functionality of chromosomal islands. **(a)** Distribution of LD block 3.1 on strain SM3. Bars above the chart represent the classification of gene groups found in the area. **(b)** Gene organization of the avhB/tra type IV secretion system from SM3. **(c)** Distribution of introgression scores for LD block 3.2. Coloured bars above the chart represent the classification of gene groups found in the area. **(d)** Illustration of synteny between gene groups in LD block 3.2 for strains lacking an insert (SM4, SM100), with the avhB/Tra conjugative system (SM3, SM121B), with a DNA rearrangement gene cluster (SM170C, SM153D), and one strain with both inserts (SM113). Dot plots above the gene group lines represent the introgression score for each gene in the gene group. Green dots represent the genes found in LD block 3.2.

### Chromosomal introgression depends on specialized transfer systems

There was a clear substructure in the LD patterns among the genes in the chromosomal cluster (**Figure 3b**, LD Block 3), and we examined the larger LD blocks in greater detail. The largest block comprised 12 genes (**Figure 3b**, LD Block 3.1), most of which were present in nearly all of the 196 strains. Cluster 3.1 included a number of hypothetical proteins, a NIPSNAP family containing protein, a phage shock protein PspA and others (**Table S9,** **Figure 4a**). We also observed toxin-antitoxin (VapC/YefM) genes (group696 and group697) in LD with this cluster (**Table S7**). However, we did not find genes that could directly explain the mobility of this introgressed region.

The second largest cluster (**Figure 3b**, LD Block 3.2) comprised six genes including a LysR family transcriptional regulator, an antibiotic biosynthesis monooxygenase, an exopolyphosphatase, TPR repeat-containing protein and an ABC transporter ATP-binding protein. To check whether the cluster could be in LD with genes that may explain its mobility, but which had not been detected by the stringently filtered introgression analysis (see Material and Methods), we extracted the genes in strongest LD with the six genes in the cluster 3.2 (**Table S7**). Three genes appeared to be in strong LD with at least one type IV secretion protein. The introgressing genes were found in different genomic contexts, and are likely chromosomal core genes that have been mobilised by different types of transfer systems (**Figure 4b-d**). In SM3 and SM121B the introgressing genes were downstream of a complete type IV secretion system, which resembles the *Agrobacterium tumefaciens* AvhB system [Chen et al., 2002] (**Figure 4b-d**). In SM170C and SM153D, another type of mobility system containing mostly hypothetical proteins along with some DNA-rearrangement genes and integrases neighbored the introgressed genes (**Figure 4d, Table S7**). In SM4 and SM100 the same core genes are present, but the transfer system has likely been lost.

### Symbiosis gene introgression is driven by a few conjugative plasmids

Symbiosis genes were in the tail of the introgression score distribution (**Figure 3a**), and a detailed analysis of three symbiosis genes (*nifB*, *nodC* and *fixT*) confirmed these patterns of HGT (**Figure 5a-c**). We also observed a complex LD pattern for the clusters comprising the symbiosis genes (**Figure 3b**, LD block 1-2), which is consistent with the presence of multiple accessory genes in distinct symbiosis plasmids within the species complex. To understand the mechanisms behind sym-gene introgression we investigated the symbiosis plasmids further. Where the assembly was complete enough to assign symbiosis genes to a specific plasmid, there was a clear pattern. Genospecies A symbiosis plasmids are all Rh06, in gsB they are Rh07, gsC has mostly Rh04 but some Rh07 and Rh08, gsD has Rh08, gsE has mostly Rh08 but some Rh06 and Rh07 (**Figure 2**; **Figure 5d; Table S5**). There are striking differences in the apparent mobility of these plasmids. Conjugal transfer genes (*tra* and *trb*) are present in some Rh06 plasmids and in all Rh07 and Rh08 plasmids, including those that are symbiosis plasmids. These transfer genes are all located together immediately upstream of the *repABC* replication and partitioning operon, in the same arrangement as in the plasmid p42a of *R. etli* CFN42, which has been classified as a Class I, Group I conjugation system [Wetzel et al., 2015]. Some *repA* sequences of sym plasmids from strains of different genospecies are identical or almost identical in sequence (**Figure 5e** **and Figure S12**). The phylogenies of the corresponding conjugal transfer genes (e.g. *traA*, *trbB* and *traG*) show the same pattern (**Figure S13**), indicating that symbiosis plasmids have crossed genospecies boundaries through conjugation. Rh08 is the most striking example (**Figure 5e**), since all strains containing a Rh08 sym-plasmid were found in an introgressed clade (**Figure S14**). We investigated the impact of this plasmid on introgression by repeating the introgression analysis in the absence of strains carrying Rh08. The mean introgression scores of all LD blocks decreased as a result of removing Rh08, but did not fully drop to background levels (**Table 1**). By randomly excluding the same number of strains and excluding them from the alignments we observed a slight decrease from 16.81 to 14.97 in the average introgression score across the 171 genes (**Table S9**). When we excluded all of the strains in the *fixT* introgressed clade (**Figure 5c, Figure S14**), which includes strains carrying Rh08 or Rh07, the introgression scores of the plasmid-borne LD blocks (**Figure 3b**, LD blocks 1 and 2) decreased greatly, whereas the chromosomal genes (**Figure 3b**, LD block 3) were less affected (**Table 1**).

**Figure 5.**
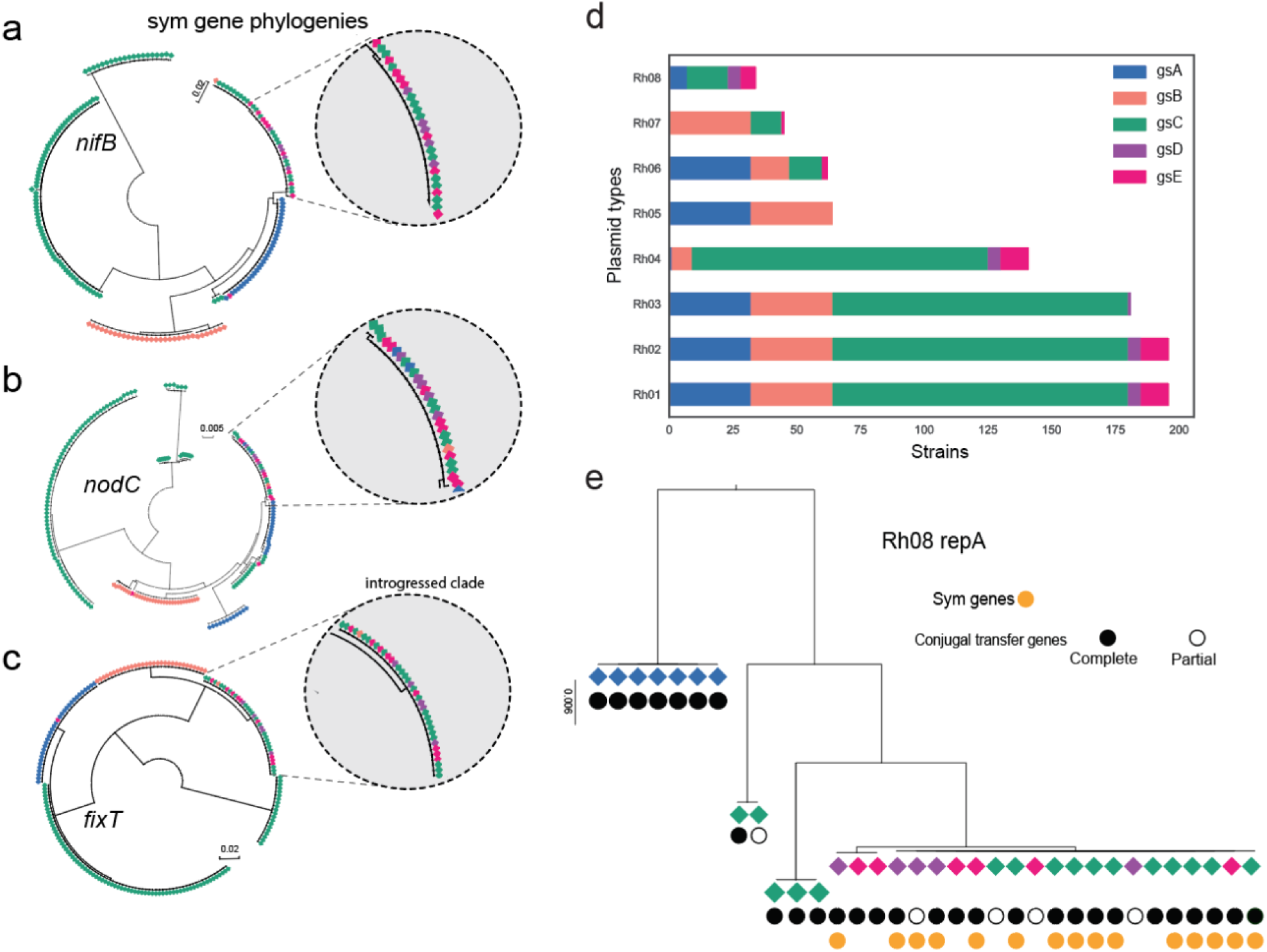
Evidence of horizontal gene transfer between genospecies. **(a)-(c)** Examples of symbiosis gene phylogenies, with insets showing clades in which identical alleles are shared across genospecies. **(d)** The distribution of plasmid groups, which were defined based on the genetic similarity of the RepA plasmid partitioning protein. **(e)** Phylogenetic analysis of the *repA* gene of plasmid type Rh08. A complete set of conjugal transfer genes has the following genes upstream of *repA*: *traI,trbBCDEJKLFGHI,traRMHBFACDG*, with the origin of transfer (*oriT*) between *traA* and *traC*. Partial sets are broken by the end of the scaffold, mostly after *traM*. The color of the diamond indicates the genospecies origin with reference to panel (d).

**Table 1.** Mean introgression score with and without introgressed clade.

| LD block | Introgression score<br>all strains | Introgression score<br>without Rh08 strains | Introgression score<br>without introgressed<br>clade |
| --- | --- | --- | --- |
| LD block 1 | 17.46 | 7.49 | 3.10 |
| LD block 2 | 17.47 | 7.65 | 3.00 |
| LD block 3 | 13.18 | 9.02 | 8.93 |
| LD block 4 | 24.00 | 13.66 | 6.00 |
| Symbiosis genes | 20.81 | 9.43 | 3.00 |

### Some *fix* genes show variation with respect to replicon location

Our LD analysis also singled out a small group of three genes that were in strong LD with each other, showed no LD with the chromosomal cluster and limited LD with the symbiosis cluster (**Figure 3b** block 4). These include *fixH*, *fixG* and a gene encoding an FNR-like protein, which are usually associated with a larger cluster of genes, *fixNOQPGHIS*, that are essential for symbiotic nitrogen fixation [Young et al. 2006]. However, they are atypical in several ways, as they have a high GC content similar to that of the core genome, they do not show the high Tajima’s D values we found typical of the main symbiosis genes, and they show variation with respect to the replicon they are associated with. In some strains, they are placed on symbiosis plasmids, in others they are located in the chromosome; other strains have two copies of the gene placed in two different genomic compartments (**Supplementary Table S5**). The introgression signal is greatly reduced when the *fixT* introgressed clade is removed (**Table 1**), implying that most of the introgression of block 4 is mediated by the mobile Rh08 and Rh07 symbiosis plasmids.

### Symbiosis genes show a unique selection signature

The chromosomal and plasmid-borne genes that exhibited introgression were not in LD and their mobility appeared to depend on different transfer systems. We wanted to investigate if the differences between the two classes of genes displaying introgression extended to selection signatures. We therefore calculated Tajima’s D, which detects deviations from the expected level of nucleotide diversity based on the number of segregating sites and pairwise differences within each gene group. Across all 196 strains, only relatively few genes showed high Tajima’s D values (**Table S5**) indicating deviations from neutral evolution. The genes within the symbiosis clusters (LD blocks 1-2) were prominent among these, making up to 57 out of the genes with the top 250 Tajima’s D scores. Since Rh08 appeared to have spread rapidly with very limited accumulation of diversity, this plasmid could be the driver of the high Tajima’s D observed for the symbiosis genes. Again, we evaluated this by excluding Rh08-bearing strains from the analysis and re-calculating Tajima’s D (**Table 2, Table S9**). We found that plasmid LD blocks (LD blocks 1 and 2) showed decreased Tajima’s D values on exclusion of Rh08 strains, while Tajima’s D values for chromosomal genes (LD block 3) were generally unaffected (**Table 2, Table S9**). We then calculated Tajima’s D values exclusively for strains found in the introgressed clade (*fixT*, **Figure 5c**), which includes both Rh08 and Rh07 carrying strains. The resulting Tajima’s D values for the symbiosis genes were negative, consistent with fewer haplotypes than expected based on the number of segregating sites (**Table 2**).

**Table 2.** Tajima’s D with and without given strains sets.

| LD block | Tajima's D all strains | Tajima's D without Rh08 strains | Tajima's D without introgressed clade | Tajima's D of only introgressed clade |
| --- | --- | --- | --- | --- |
| LD block 1 | 2.40 | 1.00 | 1.13 | -0.81 |
| LD block 2 | 1.90 | 0.66 | 0.51 | 0.08 |
| LD block 3 | 0.45 | 0.42 | 0.47 | 0.54 |
| LD block 4 | -0.15 | -0.32 | -0.001 | 0.24 |
| Symbiosis genes | 2.58 | 2.49 | 2.75 | -0.62 |

Interestingly, after excluding all Rh08 strains or the clade of introgressed strains (Rh08 and some Rh07), symbiosis genes still retained high Tajima’s D values. This indicates that multiple symbiosis gene haplotypes are also maintained at intermediate frequencies in the set of strains that does not exhibit symbiosis gene introgression. Therefore, the elevated Tajima’s D values can not be attributed solely to the existence of distinct versions of mobile symbiosis plasmids that have spread rapidly through the species complex.

Although the known symbiosis genes showed Tajima’s D patterns that were distinct from the average behavior of the genes in the plasmid-borne LD blocks, there were other genes in these blocks that showed similar patterns (**Table S8**), suggesting that they may be either under direct selection, e.g. having unknown roles in symbiosis, or might be hitchhiking with symbiosis genes under selection.

## Discussion

### Robust detection of introgression events based on gene tree traversal

HGT or introgression events in bacteria are often inferred using parametric or phylogenetic methods. Parametric methods [Lawrence and Ochman 2002; Azad and Lawrence 2007; van Passel et al., 2005] are most well suited for detecting introgression events between distantly related species, where introgression results in markedly different genomic signatures, such as abrupt changes in GC content. Detection of introgression between more closely related species, such as the members of the *R. leguminosarum* species complex described here, requires the use of phylogenetic methods that rely on gene trees derived from carefully constructed groups of orthologous genes. Because of the clear grouping of our strains into five distinct species (**Figure 1a**), we chose a simplified phylogenetic tree-traversal approach. Counting the number of transitions between genospecies on traversal proved to be a robust method for detecting introgression events, as we detected the symbiosis genes, which were candidates *a priori*. In addition, the method frequently detected groups of physically co-located and genetically linked genes, although the genes were analysed independently (**Figure 3b**). The method is mainly limited by the accuracy of the gene trees and the level of differentiation between the species for each gene group, but we found that filtering away genes with too few segregating sites was efficient in controlling the false positive rate. Another limitation is that our approach requires gene groups of a certain size, meaning that it can not be used to detect introgression of accessory genes present at low frequency within the population. Here, we limited analysis of introgression to gene groups with more than 50 members.

### Analysis of intergenic LD helps to resolve distinct introgression events

Within the *R. leguminosarum* species complex, the symbiosis genes are carried by different plasmid types (**Figure 2**), and variation in gene order and content create complex syntenic relationships (**Figure S9**). LD analysis is therefore a convenient way of understanding which introgressed genes travel together. The Mantel test has frequently been used in the comparison of genetic divergence with geographical distances [Diniz-Filho et al., 2013]. In the present study, we have applied it to calculate the genetic correlations (LD) among genes by comparing their genetic relationship matrices (GRM) [VanRaden 2008]. When autocorrelation of the GRM elements exists, possibly driven by population structure, then a relatively high false positive rate is observed [Harmon and Glor, 2010; Rousset 2002]. Aware of this effect, we used the method proposed by Mangin et al., 2012 and corrected the bias due population and phylogenetic structure. This approach is also frequently used for population structure correction in genome-wide association studies [Sauvage et al., 2014; Mamid et al., 2014]. To our knowledge, this is the first example of using a Mantel test combined with population structure-corrected pseudo-SNPs for estimation of intergenic LD. We found that calculating LD using this procedure resolved the LD inflation problem (**Figure S11**), allowing us to reliably cluster the introgressed genes based on their LD patterns.

### Introgression within the *R. leguminosarum* species complex is rare

Our introgression analysis clearly showed that genes travel across species boundaries within the species complex. Perhaps the most surprising finding was that the vast majority of genes showed no evidence of HGT, indicating that introgression events are rare. The sympatric, closely related species were thus well-separated with respect to gene flow, and specialized, conjugative transfer mechanisms appear to be required for genes to cross species barriers. We found that one of the chromosomal introgressed regions (LD block 3.2) likely represented an ICE. The *avhB* gene cassette and the *traG* gene of the type IV secretion system of this putative ICE resembles a conjugative transfer system encoded by the *virB*/*traG* of the plasmid pSymA of *S. meliloti* [Galibert et al., 2000, Barnett et al., 2001] and the *virB*/*virD4* of *Bartonella tribocorum* [Schulein et al., 2002]. However, both T4SSs in *A. tumefaciens* and *S. meliloti* (AvhB and VirB, respectively) mediate the transfer of whole plasmids, whereas we are proposing that the T4SS encoded in LD block 3.2 mediates the transfer of an integrative conjugative element (ICE). Other integrative and conjugative elements have been observed in the rhizobial genera (*Mesorhizobium loti:* [Sullivan and Ronson, 1998]; *Azorhizobium caulinodans*: [Ling et al., 2016], *Sinorhizobium*: [Zhao et al., 2017]) and in other species (*Streptococcus agalactiae*: [Rosini et al., 2006], *Bacillus subtilis*: [Merkl, 2004], *V. cholerae*: [Heidelberg et al., 2000]). Likewise, symbiosis plasmid transfer appears to require that the plasmids harbor a functional conjugal transfer system (traI,trbBCDEJKLFGHI,traRMHBFACDG), which is the case for all strains in the introgressed clade (**Figure 5, Fig S12-13-14**).

### Symbiosis gene transfer is mediated by conjugative plasmids

The occurrence of HGT of symbiosis genes within and between distant rhizobial genera (*Rhizobium*, *Bradyrhizobium*, *Sinorhizobium*, *Azorhizobium*, and *Mesorhizobium*), nodulating different legume species, has been widely reported [Pérez-Carrascal et al., 2016; Hirsch et al, 1980; Rogel et al., 2011; Lemaire et al., 2015; Andrews et al., 2018]. This shows that symbiosis gene transfer is not restricted by genetic divergence and in many cases is not species specific [Provorov and Andronov, 2017; Greenlon et al., 2019].

Here, we have shown that species-specific clades still exist even among symbiosis genes (**Figure 5a-c**). In most species-specific clades, the genes were carried on a non-mobile symbiosis plasmid (Rh04) (**Fig S14**), suggesting that, in this species complex, symbiosis gene introgression was only observed when the strain had a plasmid with a conjugation apparatus. We verified this by characterizing the plasmid diversity within the strain pool. Symbiosis plasmids belong to a number of plasmid types (Rh04, Rh06, Rh07 and Rh08), and phylogenetic evidence indicated that some of them (Rh07 and Rh08) have been transferred through conjugation between different genospecies (**Figure 5e, Figure S12**). These transfers are likely recent since many of the sequences (*repA* and *tra* genes) have not yet diverged. Because conjugation requires cell-to-cell contact, plasmid transfer is not just constrained by genetic similarity [Silva et al., 2003; Pérez-Carrascal et al., 2016], but also by the requirement that the donor and recipient are found at the same location.

### There is introgression of *fix* genes that vary in genomic location

The genes that displayed introgression and were on symbiosis plasmids (LD blocks 1 and 2) were not in LD with the introgressing chromosomal genes (LD block 3) (**Figure 3**) and they displayed different selection signatures (**Table 2**), indicating that chromosomal and plasmid-associated introgression events are independent. LD block 4 was atypical, because it contained putative symbiosis genes that showed variation with respect to replicon location and were conspicuously absent from the immobile symbiosis plasmid Rh04 (**Table S5**). These genes are part of the *fixNOQPGHIS* cluster, and it is known that this set of genes is essential for symbiotic nitrogen fixation, but that a single copy is sufficient [Young et al., 2006]. Nevertheless, their high GC content and frequent chromosomal location indicates that these are core genes that have been co-opted into a symbiotic role. Consistently, they show introgression when a copy has been acquired by one of the mobile types of symbiosis plasmid. This suggests that they have been mobilized as a consequence of their symbiotic function, perhaps because they confer an advantage when transferred to a recipient that does not have an optimal *fixNOQPGHIS* cluster for symbiosis.

### Intermediate frequency symbiosis gene haplotypes co-exist in sympatry

Just as the five genospecies co-exist, so do different symbiosis gene haplotypes and plasmids. The symbiosis genes had strikingly high Tajima’s D values, indicating an excess of intermediate-frequency haplotypes. The *fixT* gene is the gene with the fewest haplotypes, presenting only five haplotypes in total (**Supplementary Figure S14**). Four of these are present in the Danish organic fields, and the haplotype characteristic of the introgressed clade was found at trial sites in Denmark, France, and the UK as well as in Danish organic fields (**Supplementary Table S5**).

The presence of distinct groups of haplotypes at intermediate frequency could be a result of negative frequency dependent selection [Amarger & Lobreau, 1982; Provorov and Vorobyov, 2000b; Provorov and Vorobyov, 2006; Bever, 1999]. This type of balancing selection could actively maintain symbiont diversity by increasing the fitness advantage of strains when they are rare. An alternative, not necessarily mutually exclusive hypothesis, is that distinct symbiosis haplotypes are maintained by host specialization. If the selective optimum between rhizobium and its host changes over time, symbiosis gene alleles that contribute to the interaction will experience repeated partial sweeps, increasing the frequency of different adaptive alleles in different parts of the allelic range. Under balancing selection, these partial local sweeps can create elevated differentiation among allelic haplotypes and reduce nucleotide and haplotype diversity in the regions flanking each selected locus [Yoder et al., 2014; Garud et al., 2015].

The 196 strains characterized here were all collected from clover root nodules, and the colonisation of nodules is a bottleneck that imposes strong selection. We see that certain haplotypes of symbiosis-related genes have introgressed across multiple genospecies, implying that these genes provide a fitness benefit that is largely independent of the genomic background. However, this pattern of selection appears to be exceptional, because the number of other genes that showed a similarly high introgression signal was very limited. Most of the thousands of accessory genes in the gene pool are not strongly introgressing, suggesting that they are contributing to the adaptive differences that presumably distinguish the different genospecies. Judging from our results, the high mobility of symbiosis genes, extensively documented in the literature, is not typical of the accessory genome in general.

## Conclusions

Using new methods for detection of introgression events and intergenic LD analysis, we carried out an unbiased investigation of introgression within an *R. leguminosarum* species complex. We found that introgression was generally very limited, with most genes displaying genetically distinct, species-specific variants. Striking exceptions are the genes located on symbiosis plasmids, especially the symbiosis genes, and a limited number of chromosomal islands, which appear to travel across species boundaries using conjugative transfer systems. The plasmid and chromosomal introgression events are independent and subject to different selective pressures, and some genes appear to move both between species and between replicons.

## Material and Methods

### Rhizobium sampling and isolation

White clover (*Trifolium repens*) roots were collected from three breeding trial sites in the United Kingdom (UK), Denmark (DK), and France (F) (Figure S1A), and 50 Danish organic fields (DKO) (Figure S1b). Roots were sampled from 40 plots from each trial site. The total number of plots was 170. The samples were stored at ambient temperature for 1-2 days and in the cold room (2) for 2-5 days prior to processing. Pink nodules were collected from all samples, and a single bacterial strain was isolated from each nodule as described by [Bailly et al., 2011]. From each plot, 1 to 4 independent isolates were sampled. In total 249 strains were isolated from *T. repens* nodules. For each site the clover varieties were known, and representative soil samples from clover-free patches were collected and sent for chemical analysis. Furthermore, latitude and longitude data were collected (**Table S1**).

### Genome assembly

A set of 196 strains was subjected to whole genome shotgun sequencing using 2×250 bp Illumina (Illumina, Inc., USA) paired-end reads by MicrobesNG (https://microbesng.uk/, IMI - School of Biosciences, University of Birmingham). In addition, 8 out of the 196 strains were re-sequenced using PacBio (Pacific Biosciences of California, Inc., USA) sequencing technology (Table S2, Figure S2). Analysis of 16S rDNA confirmed that all 196 of the strains were *Rhizobium leguminosarum*.

Genomes were assembled using SPAdes (v. 3.6.2) [Bankevich et al., 2012]. SPAdes contigs were cleaned and assembled further, one strain at a time, using a custom Python script (Jigome, available at https://github.com/jpwyoung/genomics). First, low-coverage contigs were discarded because they were mostly contaminants from other genomes sequenced in the same Illumina run. The criterion for exclusion was a SPAdes k-mer coverage less than 30% of the median coverage of putative single-copy contigs (those *>* 10kb). Next, putative chromosomal contigs were identified by the presence of conserved genes that represent the syntenic chromosomal backbone common to all *R. leguminosarum* genospecies. A list of 3215 genes that were present, in the same order, in the chromosomal unitigs of all eight of the PacBio assemblies was used to query the Illumina assemblies using *blastn* (≥90% identity over ≥90% of the query length). In addition, contigs carrying *repABC* plasmid replication genes were identified using a set of *RepA* protein sequences representing the twenty distinct plasmid groups found in these genomes (*tblastn* search requiring ≥95% identity over ≥90% of the query length). A ‘contig graph’ of possible links between neighbouring contigs was created by identifying overlaps of complete sequence identity between the ends of contigs. The overlaps created by SPAdes were usually 127 nt, although overlaps down to 91 nt were accepted. Contigs were flagged as ‘unique’ if they had no more than one connection at either end, or if they were *>* 10 kb in length. Other contigs were treated as potential repeats. The final source of information used for scaffolding by Jigome was a reference set of *R. leguminosarum* genome assemblies that included the eight PacBio assemblies and 39 genomes publicly available in GenBank. A 500-nt tag near each end of each contig, excluding the terminal overlap, was used to search this database by blastn; high-scoring matches to the same reference sequence, with the correct spacing and orientation, were subsequently used to choose the most probable connections through repeat contigs. Scaffolding was initiated by placing all the chromosomal backbone contigs in the correct order and orientation, based on the conserved genes that they carried, and extending each of them in both directions, using the contig graph and the pool of remaining non-plasmid contigs, until the next backbone contig was reached or no unambiguous extension was possible. Then each contig carrying an identified plasmid origin was similarly extended as far as possible until the scaffold became circular or no further extension was justified, and unique contigs that remained unconnected to chromosomal or plasmid scaffolds were extended. Finally, scaffolds were connected if their ends had appropriately spaced matches in the reference genomes. Scaffold sequences were assembled using overlap sequences to splice adjacent contigs exactly, or inserting an arbitrary spacer of twenty “N” symbols if adjacent contigs did not overlap. The *dnaA* gene (which was the first gene in the chromosomal backbone set and is normally close to the chromosomal origin of replication) was located in the first chromosomal scaffold, and this scaffold was split in two, with chromosome-01 starting 127 nt upstream of the ATG of *dnaA* and chromosome-00 ending immediately before the ATG. The remaining chromosomal scaffolds were numbered consecutively, corresponding to their position in the chromosome. Plasmid scaffolds were labelled with the identifier of the *repA* gene that they carried. Scaffolds that could not be assigned to the chromosome or a specific plasmid were labelled ‘fragment’ and numbered in order of decreasing size. Subsequent analysis revealed large exact repeats in a few assemblies. These were either internal inverted repeats in the contigs created by SPAdes (5 instances) or large contigs used more than once in Jigome assemblies (18 instances). They were presumed to be artifacts and removed individually. Assembly statistics were generated with QUAST (v 4.6.3, default parameters) [Gurevich et al, 2013]. (Figure S3). Genes were predicted using PROKKA (v 1.12) [Seemann, 2014]. In summary, genomes were assembled into [10-96] scaffolds, with total lengths of [8355366-6967649] containing [6,642-8,074] annotated genes, indicating that we have produced assemblies of reasonable quality, which comprehensively captured the gene content of the sequenced strains (Table S2 and S3).

### Orthologous genes prediction

Orthologous gene groups were identified among a total of 1,468,264 predicted coding sequences present across all (196) strains. We used two software packages for ortholog identification: Proteinortho [Lechner et al., 2014] and Syntenizer3000 (https://github.com/kamiboy/Syntenizer3000/). The software Proteinortho (v5.16b), was executed with default parameters and the synteny flag enabled, to predict homologous genes while taking into account their physical location. For the analysis in this paper, we were only interested in orthologs and not paralogs. Paralogous genes predicted by Proteinortho were filtered out by analyzing the synteny of homologous genes surrounded by a 40-gene neighbourhood (see Synteny section). After this filtering step, the orthologous gene groups were aligned using ClustalO ([Sievers et al., 2011], v. 1.2.0). Each gene sequence was translated into its corresponding amino acid sequence before alignment and back-translated to the original nucleotides. Each gap was replaced by 3 gaps, resulting in a codon-aware nucleotide alignment.

### Synteny

First, gene groups were aligned with their neighbourhoods (20 genes each side) using a modified version of the Needleman-Wunsch algorithm [Needleman and Wunsch, 1970]. We counted the number of gene neighbours that were syntenic across strains before a collinearity break. We used this score to disambiguate gene groups that contain paralogs. Paralogs are the result of gene duplication, and as such one of the paralogs is the original, and the rest are copies. Based on similarity, we kept the least divergent gene inside of the original homology group while removing the copied paralogs, if possible into a new gene group designated group name “-”2. Orphan genes that were present only in one strain, were removed from the analysis.

### Variant Calling

Codon-aware alignments were used in order to detect single nucleotide polymorphisms (SNPs). For a given gene alignment and position, we first counted the number of unique nucleotides (A, C, T, G). Sites containing 2 unique nucleotides were considered variable sites (bi-allelic SNPs). After finding variable sites, SNP matrices were encoded as follows: major alleles were encoded as 1 and minor alleles as 0. Gaps were replaced by the site mean. Later steps were executed in order to filter out unreliable SNPs. We restricted the analyses to genes found in at least 100 strains. By looking at the variants and their codon context, we excluded SNPs placed in codons containing gaps, or containing more than one SNP, or with multi-allelic SNPs. Based on these criteria we ended up with 6,529 out of 22,115 genes and 441,287 SNPs. Scripts and pipelines are available at a GitHub repository (https://github.com/izabelcavassim/Rhizobium_analysis/).

### Plasmid replicon groups

Plasmid replication genes (repABC operons) were located in the genome assemblies by *tblastn*, initially using the RepA protein sequences of the reference strain 3841 as queries (Young et al., 2006). Hits covering ≥70% of the query length were accepted as repA genes, and those with ≥90% amino acid identity were considered to belong to the same replication group (putative plasmid compatibility group). Hits with lower identity were used to define reference sequences for additional groups, using sequences from published *Rhizobium* genomes when available, or from strains in this study. Groups were numbered (Rh01, etc) in order of decreasing abundance in the genome set. RepB and RepC sequences corresponding to the same operons as the RepA references were used to check whether the full *repABC* operon was present at each location, requiring ≥85% amino acid identity.

### Presence of symbiosis genes in all strains

Since all sequenced strains were isolated from white clover nodules, they are expected to carry the canonical symbiosis genes. One strain, SM168B, carried no symbiosis genes. Subsequent nodulation tests showed that the strain could colonize white clover and produce pink nodules, suggesting that the genes were lost during the pre-sequencing processing. On the other hand, strains SM165B and SM95 were found to have duplicated symbiosis regions.

### Population genetic analysis

Population genetic parameters (Tajima’s D, nucleotide diversity, average pairwise differences and number of segregating sites) were estimated using the python library dendropy [Sukumaran, 2010].

### Introgression Score

Despite the clear grouping of the 196 strains into distinct species, there was still extensive cross-species sequence conservation, allowing the construction of high-quality orthologous gene groups (**Table S4**). We took advantage of these for detecting introgression events by generating and traversing gene trees for each of the gene groups. Individual gene trees were first constructed using the neighbor-joining clustering method (software RapidNJ version 2.3.2) [Simonsen and Pedersen 2011]. Each tree was traversed based on depth first traversal algorithm [Tarjan, 1972] by visiting each node after visiting its left child and before visiting its right child, searching deeper in the tree whenever possible. When the leaf of the tree was reached, the strain number and its genospecies origin were extracted. A list containing the genospecies was stored for the entire tree. The introgression score was computed as following:

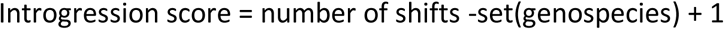

The introgression score evaluates the number of times a shift (from one genospecies to another) is observed in a branch. The minimum possible is the total number of genospecies -1 shifts. A tree congruent to the species tree would have a introgression score equal to zero (**Figure S8**).

### Intergenic Linkage Disequilibrium corrected for population structure

Sample structure or relatedness between genotyped individuals leads to biased estimates of linkage disequilibrium (LD) and increase of type I error. In order to correct for the autocorrelation present in this data, the genotype matrix *X*(coded as 0’s and 1’s) was adjusted as exemplified in Mangin et al. 2012 and Long et al. 2013.

The covariance 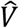 between individuals was calculated as follows:

Let *N* denote the total number of individuals and *M* the total number of markers, the full genotype matrix (*X*) has *NxM* dimensions with genotypes encoded as 0’s and 1’s. For simplicity, each SNP information is looked as vectors, *S*_(*j*,*i*)_ = 1,…, *M*.

The first step of the calculations was to apply a Z-score normalization on the SNP vectors by subtracting each vector by its mean and divide it by its standard deviation 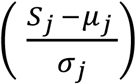.

We then computed the covariance matrix between individuals as follows.

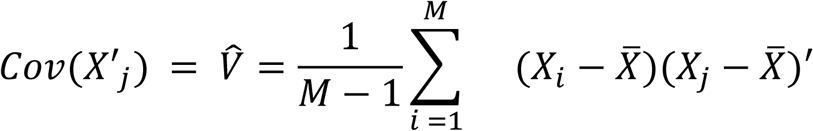

*Cov*(*X*), can also be computed by the dot product of the genotype matrix:

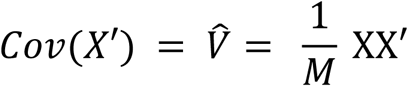

The result is an *N x M* matrix, where *N* is the number of strains. This matrix is also known as Genomic Relationship Matrix (GRM) [VanRaden, 2008]. We then decomposed the GRM matrix using linalg function of scipy (python library).

Then the ‘decorrelation’ of genotype matrix *X* was done by multiplying *X* by the inverse of the square root of 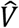 as follow:

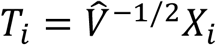

*T* is therefore the pseudo SNP matrix, which is corrected for population structure.

The correlation between genes matrices was obtained by applying mantel test on the GRM (genetic distances) between pairs of genes:

For a data set composed of a distance matrix of gene X (*D_ij_^x^*) and a genetic distance matrix of gene Y (*D_ij_^y^*), it was computed the scalar product of these matrices adjusted by the means and variances (*var*(*X*) *and var*(*y*)) of the matrices X and Y:

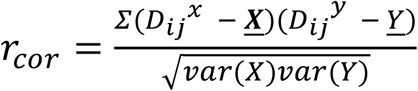

The standardized Mantel test is actually the Pearson correlation between the elements of genes X and Y.

### Filtering criteria for top introgressed genes

In order to identify genes that had trustable signals of introgression we used a stringent filtering criteria as follows: number of sequences > 50; number of segregating sites > 10; average pairwise differences > 10, ANI > 0.7, introgression score > 10.

## Author’s contributions

Conceptualization: MIAC, JPWY, SM, MHS and SUA; Methodology: MIAC, JPWY and SM; Software: MIAC, AB, BV, JPWY and CM; Validation: MIAC, CM, SM, JPWY; Formal Analysis: MIAC, JPWY, CM, SM, AB, BV and BF; Investigation: SM; Resources: SUA, JPWY and MHS; Data Curation: MIAC, CM, JPWY, SM, SUA and MHS; Writing - Original Draft: MIAC; Writing - Review and Editing: MIAC, JPWY, SUA, MHS, SM, BV; Visualization: MIAC, SM, JPWY; Supervision: SUA, JPWY, MHS; Project Administration: SUA; Funding Acquisition: SUA.

## Acknowledgements

This work was funded by grant no. 4105-00007A from Innovation Fund Denmark (S.U.A.). Genome sequencing was provided by MicrobesNG, which is supported by the BBSRC (grant number BB/L024209/1). The authors would also like to thank industrial partners DLF Trifolium, SEGES and Legume Technology Ltd. for their contribution to the field trials.

## Availability of data and materials

The data that support the findings of this study are available in the INSDC databases under Study/BioProject ID PRJNA510726. Accessions numbers are from SAMN10617942 to SAMN10618137 consecutively and are also provided in the **Supplementary table S10**. Gene alignments SNP data and metadata can be downloaded from the following folder: https://www.dropbox.com/sh/6fceqmwfa3p3fm6/AAAkFIRCf7ZxgO1a4fHv3FeOa?dl=0

## Additional Files

### Supplementary tables

This file is a multi-page table composed of the following information:

**Table S1.** Metadata: information on field trials for each isolate;

**Table S2.** Genome statistics: information on genome assemblies;

**Table S3.** Conserved genes: list of conserved genes used for species tree construction;

**Table S4.** RepA types: representatives of repA types; Rh classification and nucleotide sequences;

**Table S5.** Genes statistics: information on genes and plasmid types for each isolate;

**Table S6.** Population genetic parameters: of every orthologous gene and introgression scores;

**Table S7.** Inserts description: LD analysis between chromosomal introgressed clade and avhB description;

**Table S8.** Symbiosis genes parameters: pop. gen. parameters of symbiosis genes in contrast to recA and rpoB;

**Table S9.** Stats on the top 171 introgressed genes. Tajima’s D and introgression score stats with and without specific sets of strains;

**Table S10.** Accession numbers of the 196 genomes.

## Supplementary figures

**Figure S1:**
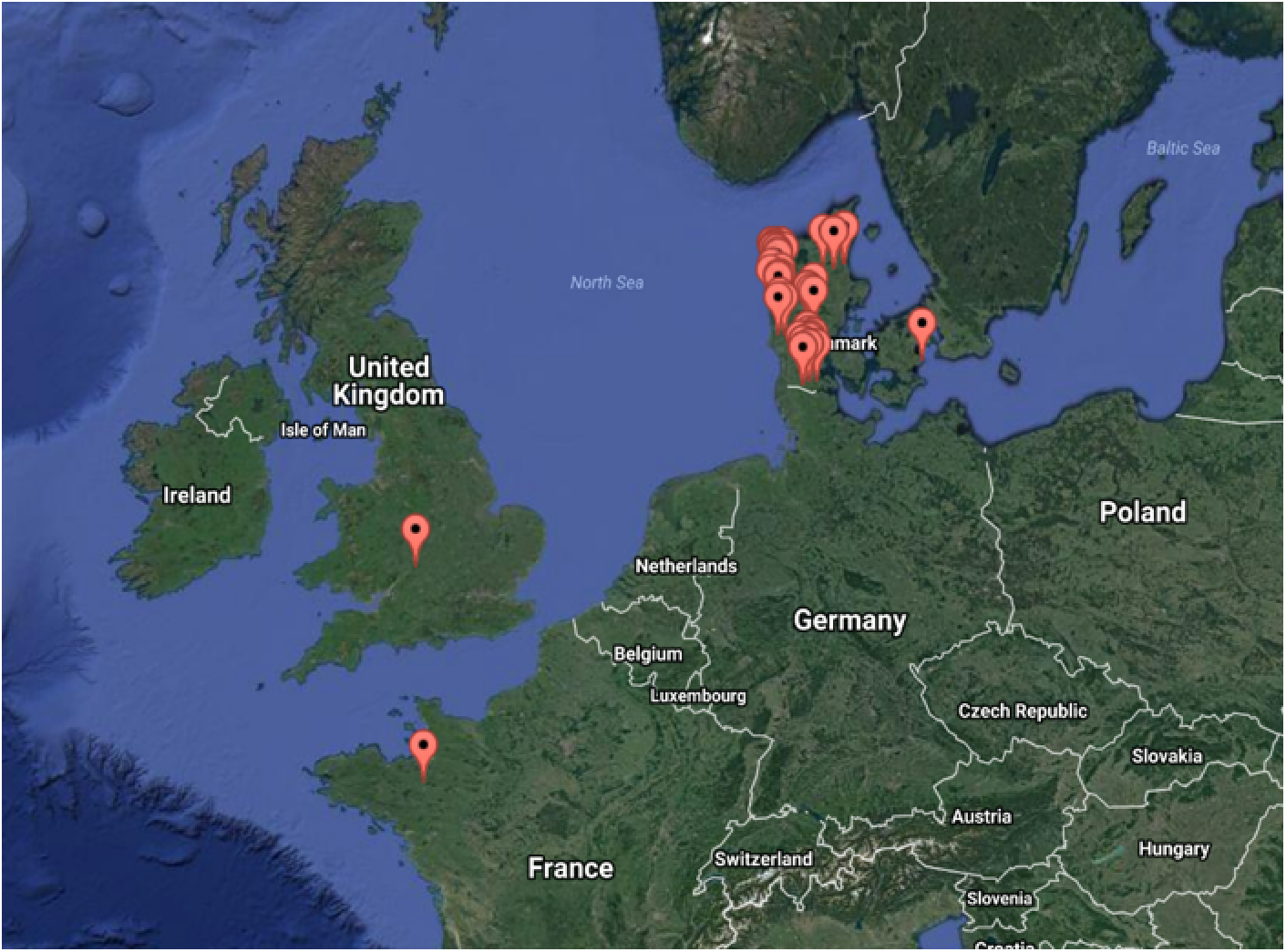
White clover roots were collected from three different DLF trials sites: United Kingdom (UK), Denmark (DK) and France (F).

**Figure S2:**
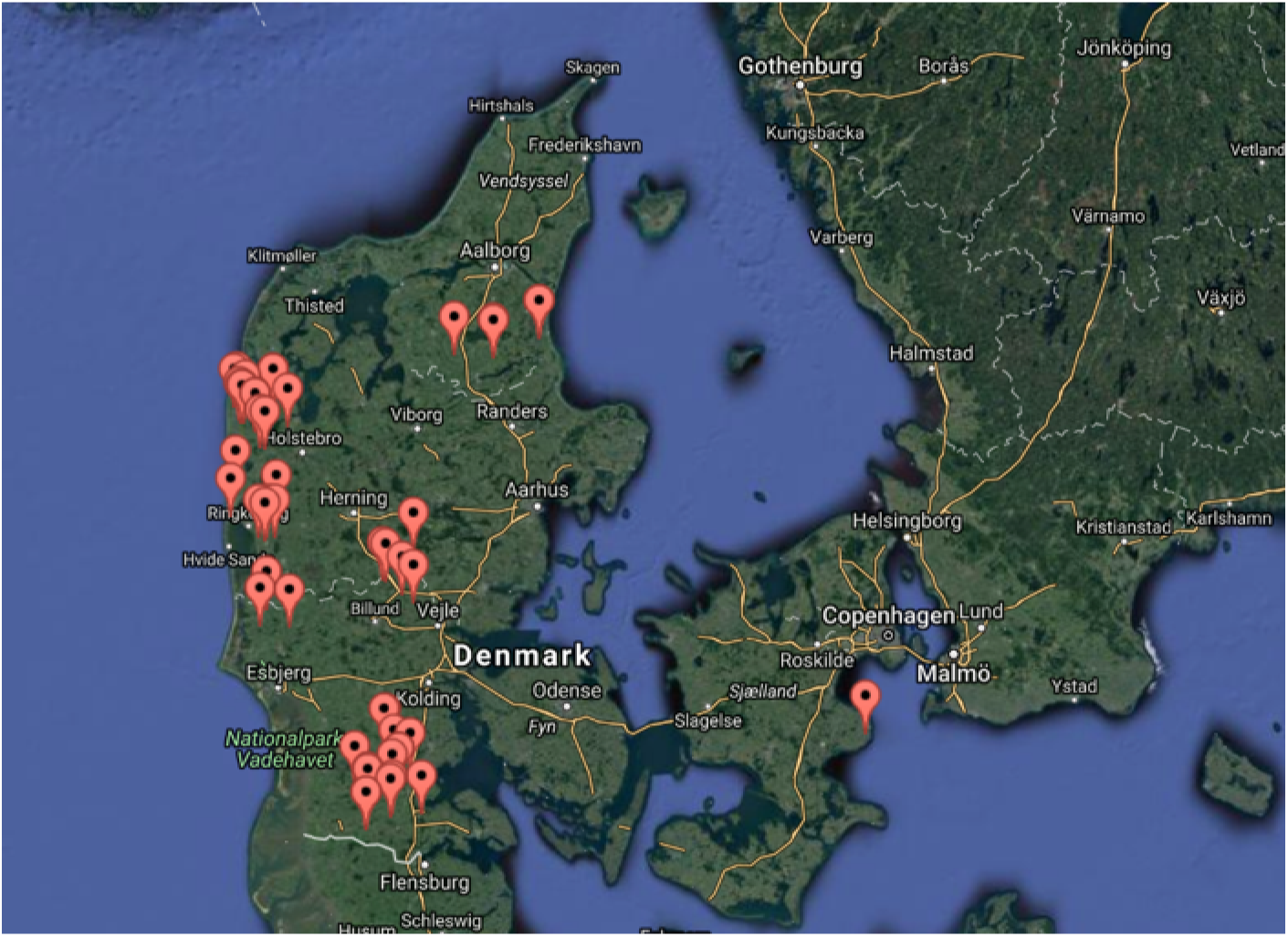
Soil samples were also collected from 50 Danish organic fields (DKO). Geographic information system (SIS) data is attached in supplementary table 1.

**Figure S3:**
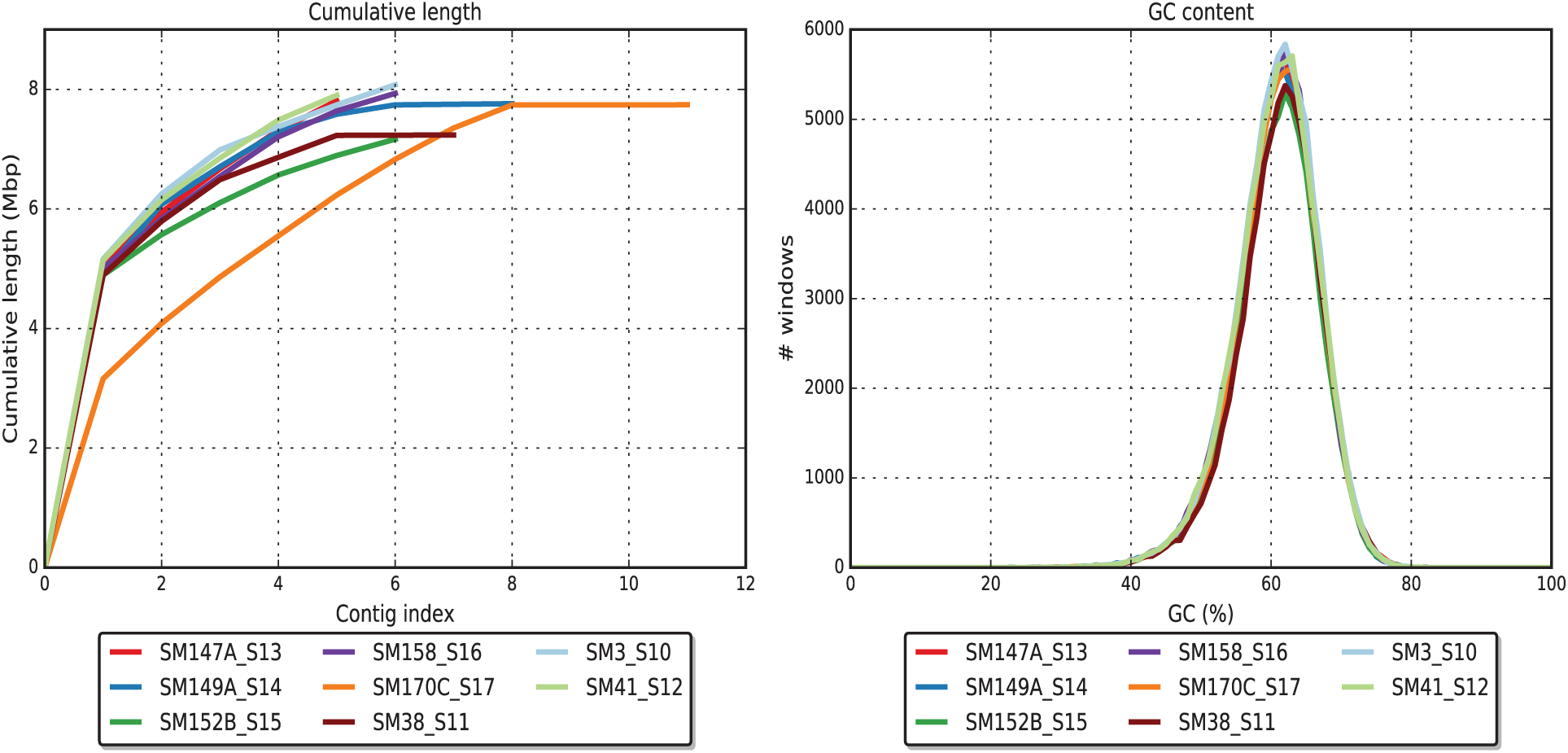
Number of contigs and GC content in each pacbio assembly. These strains were used in order to improve the illumina assemblies. Strain SM170C was excluded from the re-assembly analysis.

**Figure S4:**
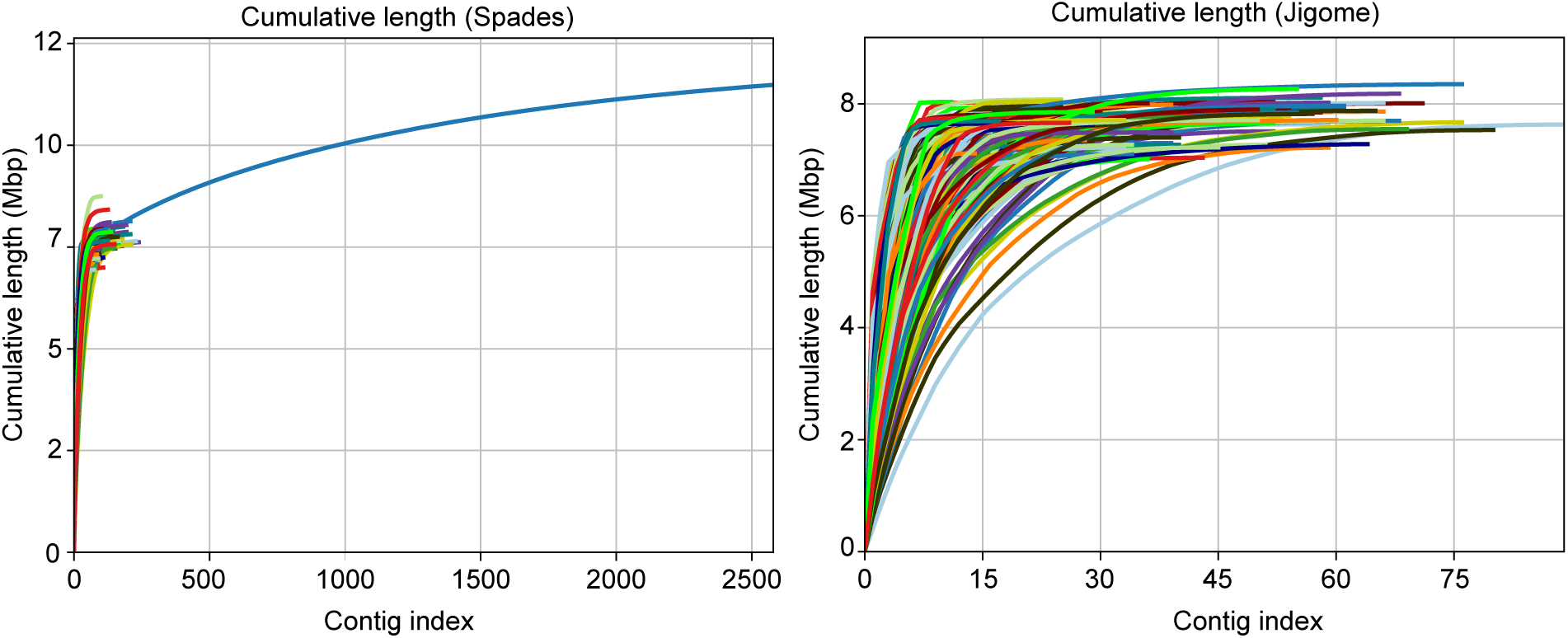
Number of contigs per strain using Spades and later Jigome. A fixed threshold for a minimum contig length of 200 bp was used.

**Figure S5:**
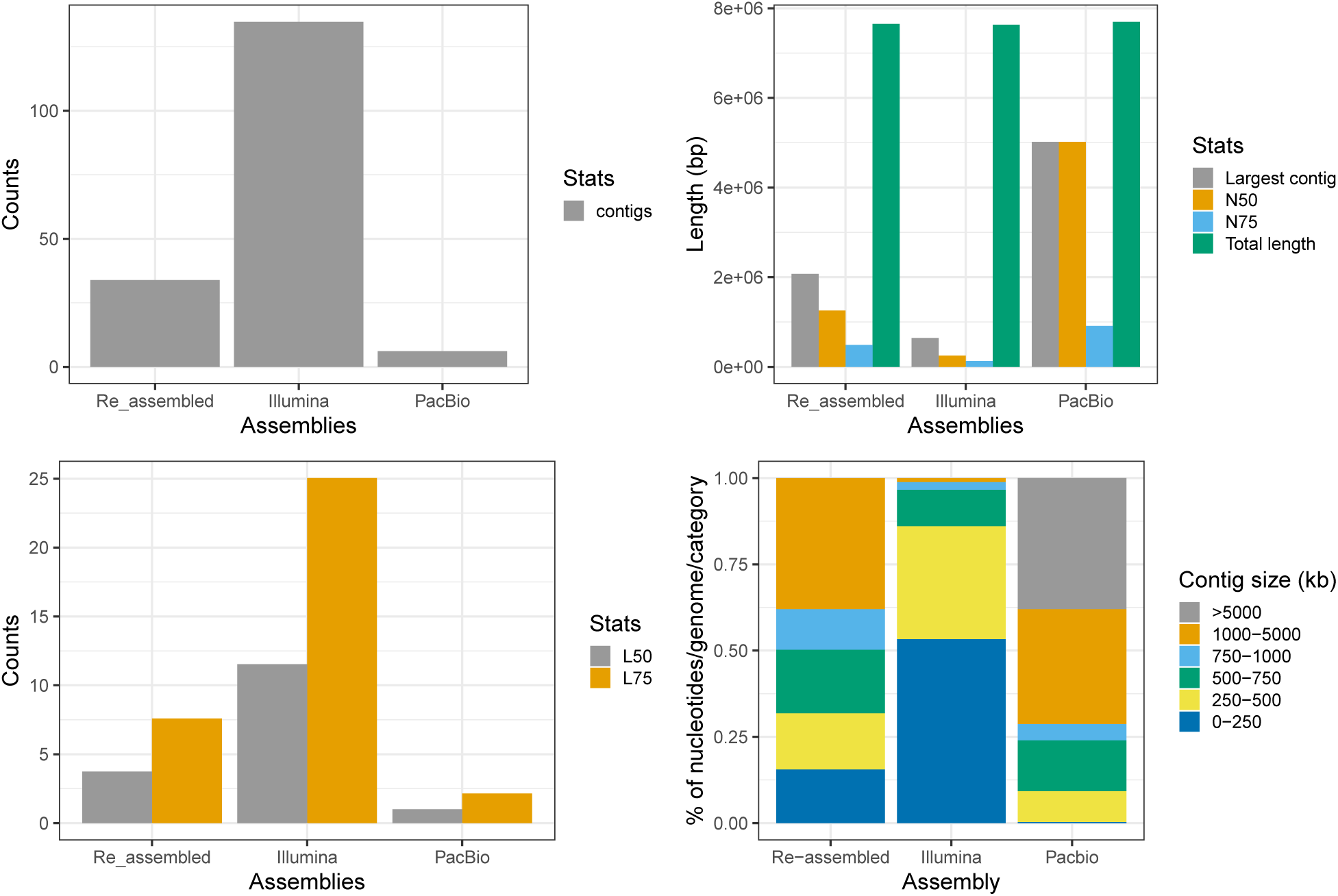
Different statistics across the 3 assemblies: Illumina (Spades assembly), Pacbio (HGAP.3 assembly) and Re-assembled (Illumina re-assembled with Jigome). Re-assembled and Pacbio were used in these analysis.

**Figure S6:**
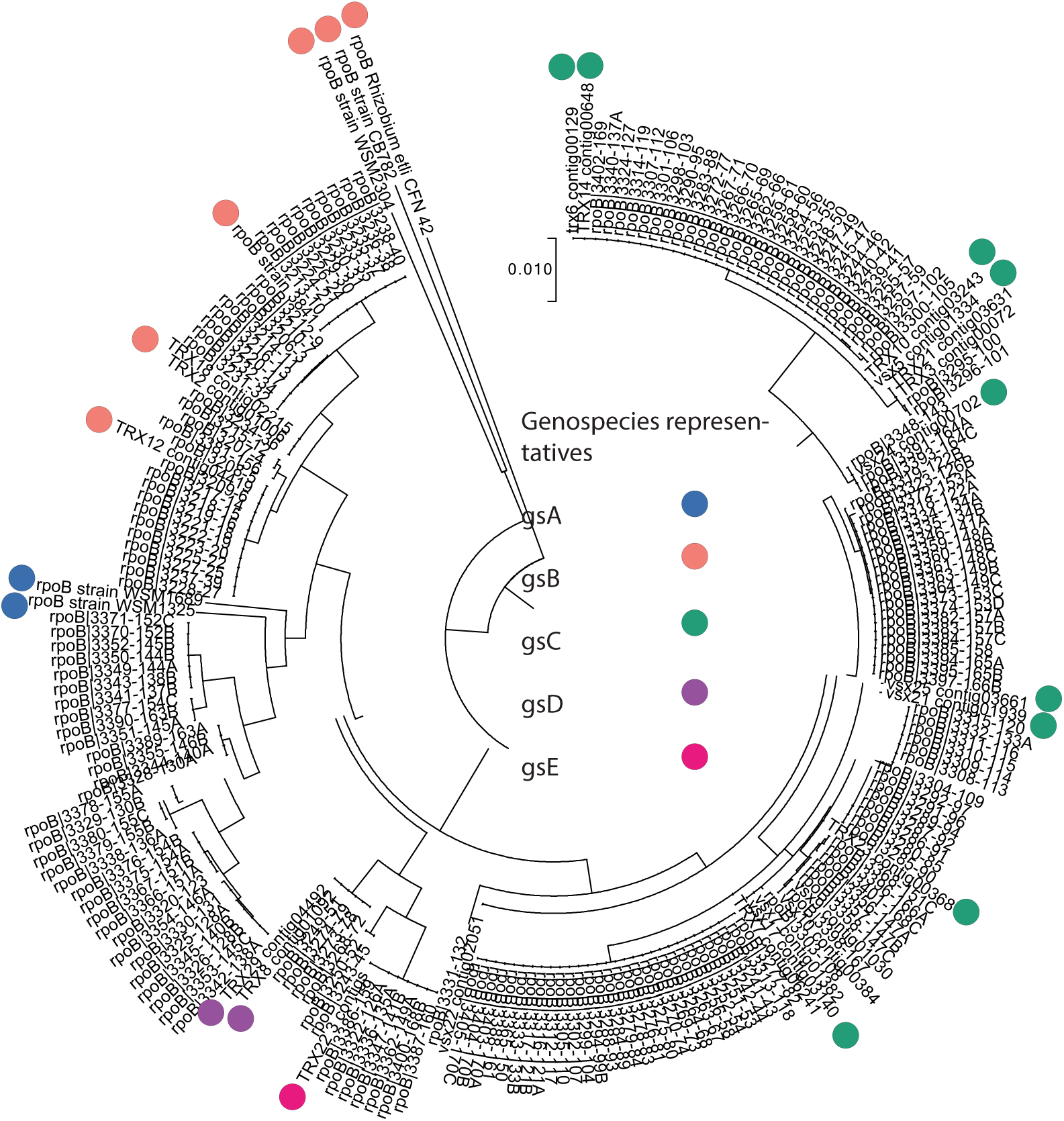
(a) RpoB phylogenetic tree and *rpoB* sequences of representatives of each genospecies (circles). These sequences were previously classified by Kumar et al., 2015.

**Figure S7:**
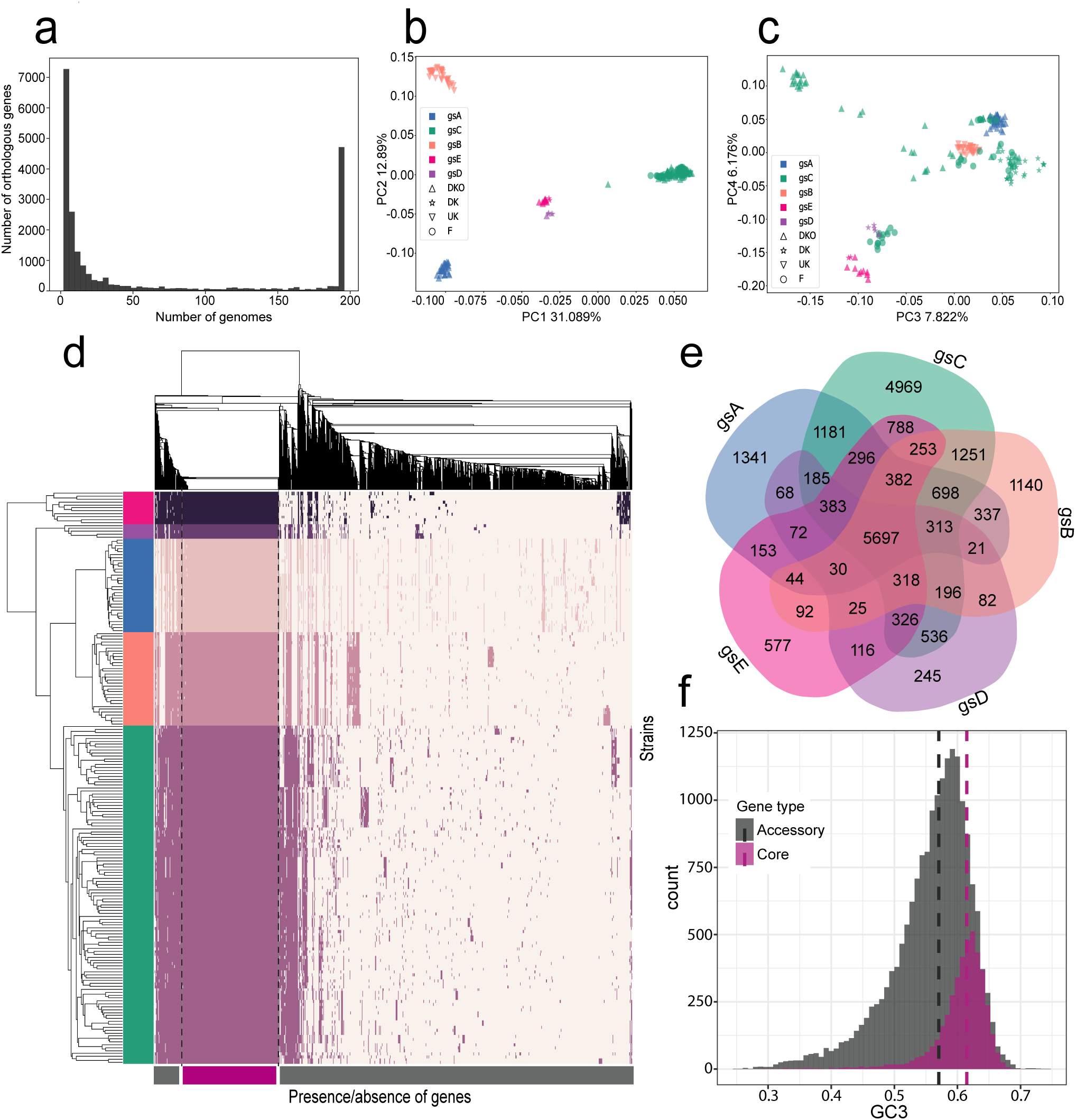
(a) Histogram showing the distribution of shared genes across strains, with a total of 22,115 orthologous genes. (b) Principal component analysis (PCA) of the covariance matrix based on the allelic variation of 6,529 genes that were present in at least 100 strains (see Methods). The colours correspond to the genospecies and the shapes to the origin of the sample. PC1 and PC2. (c) PC3 and PC4 of the PCA. (d) Matrix of the presence (dark) and absence (light) of all 22,115 orthologous gene groups. Strains are clustered by similarity (y-axis), and genes are clustered by their patterns of presence and absence (y-axis). (e) Venn diagram of the shared orthologous genes across the 5 genospecies; the outermost numbers represent the number of genes that are private to the genospecies. (f) GC3 content distribution across accessory and core genes; dashed lines represent the median GC3 of each category.

**Figure S8:**
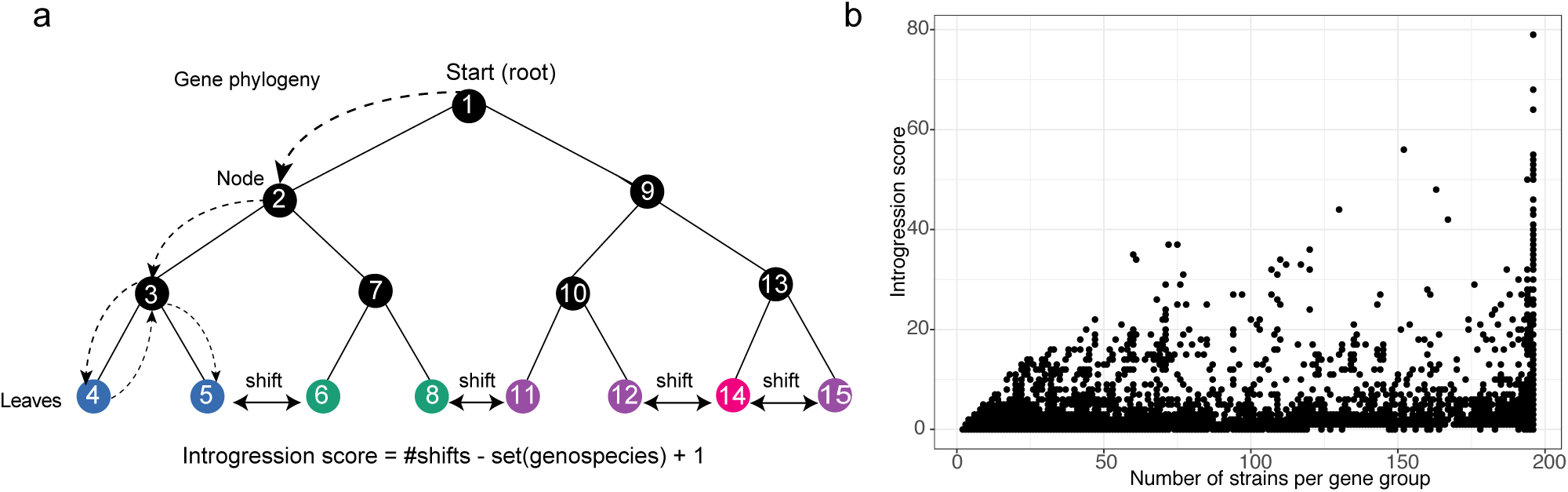
Illustration of the approach for detecting gene introgression (a), and its dependency on the number of members in each orthologous gene (b).

**Figure S9:**
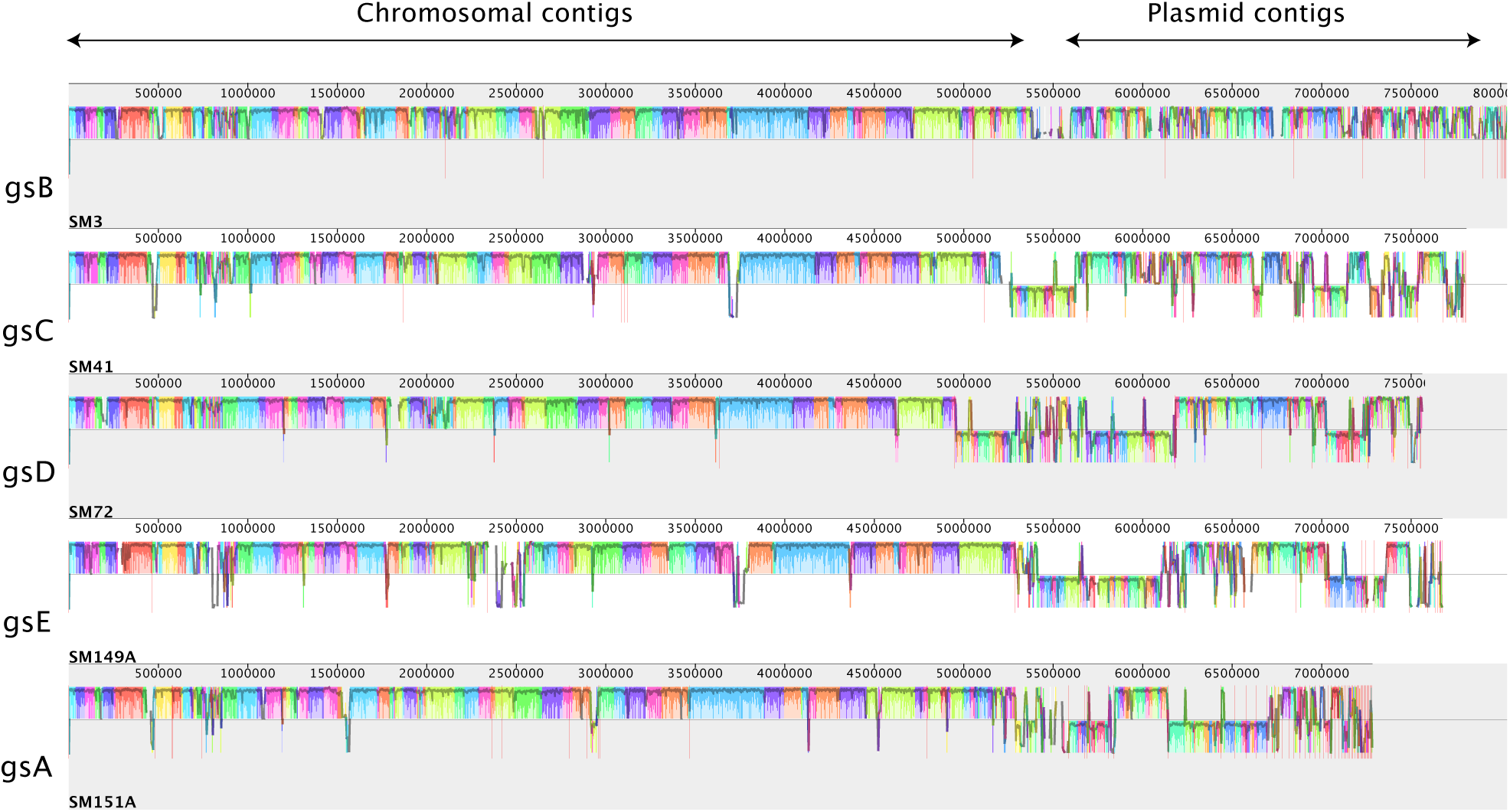
Structural rearrangements and gene interactions of *Rhizobium leguminosarum* bv. *trifolii*. High chromosomal collinearity and distribution of plasmid types. Multiple alignment across one strain from each genospecies, plasmids and chromosomal contigs are distinguished. The coloured blocks correspond to local collinear blocks that are detected by Mauve alignment and are internally free from genome rearrangements.

**Figure S10:**
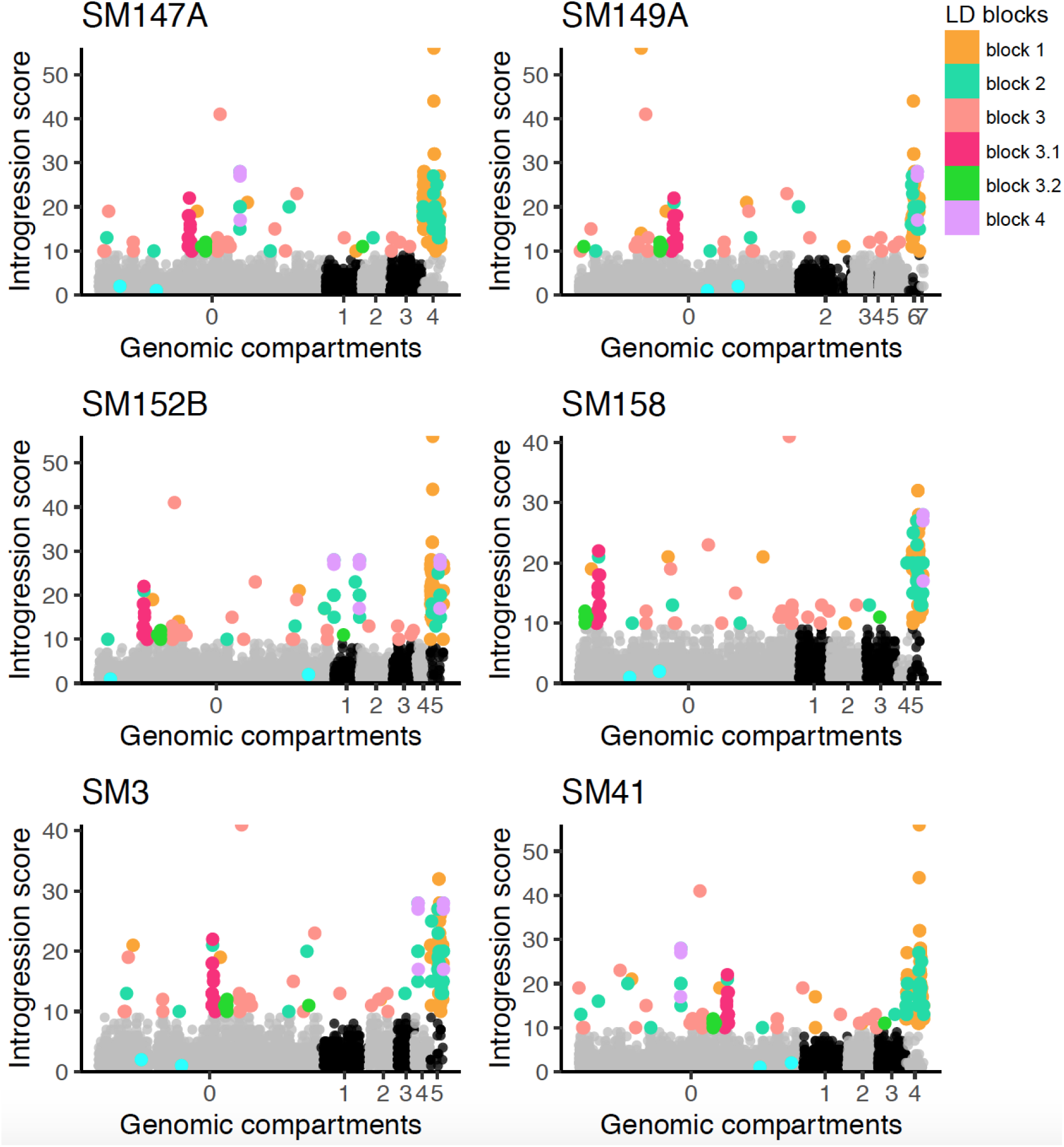
Orthologous gene groups were blast against pacbio assemblies and introgression score (y-axis) is plotted against genomic positions (x-axis). Grey and black dots represents the genes distributed in the different compartments (chromosome = 0, chromid = 1 and 2, >2 = plasmids). Light blue are the two conserved genes (*recA* and *rpooB*), all the other colors correspond to the linkage blocks classified by the intergenic LD analysis.

**Figure S11:**
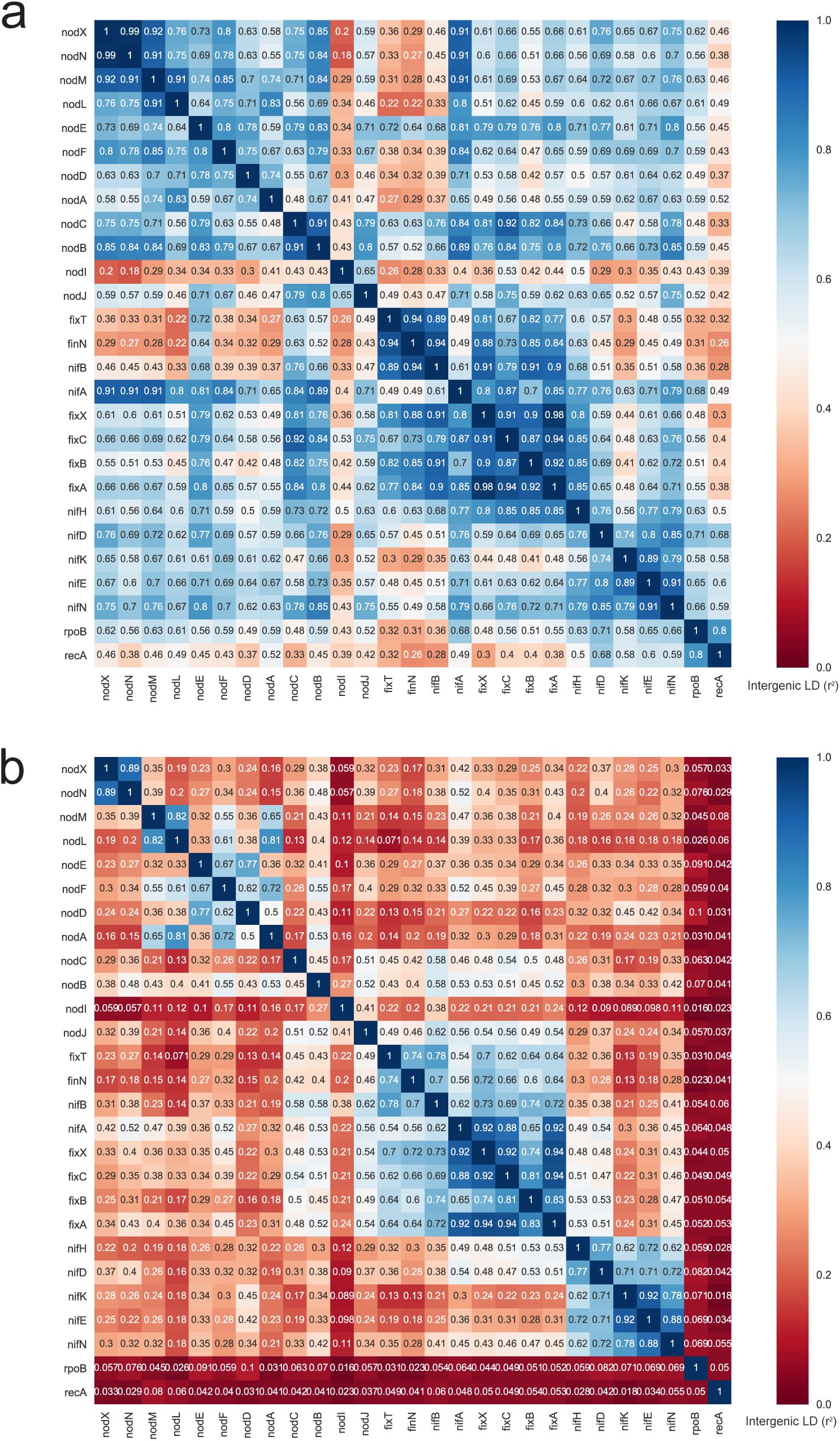
Intergenic linkage disequilibrium before (a) and after (b) population structure correction. The top 24 genes displayed in the matrix are plasmid-borne symbiosis genes, the two last genes (*rpoB* and *recA*), are highly conserved chromosomal genes: part of the DNA recombination and repair system; and part of beta subunit in RNA polymerase, respectively.

**Figure S12:**
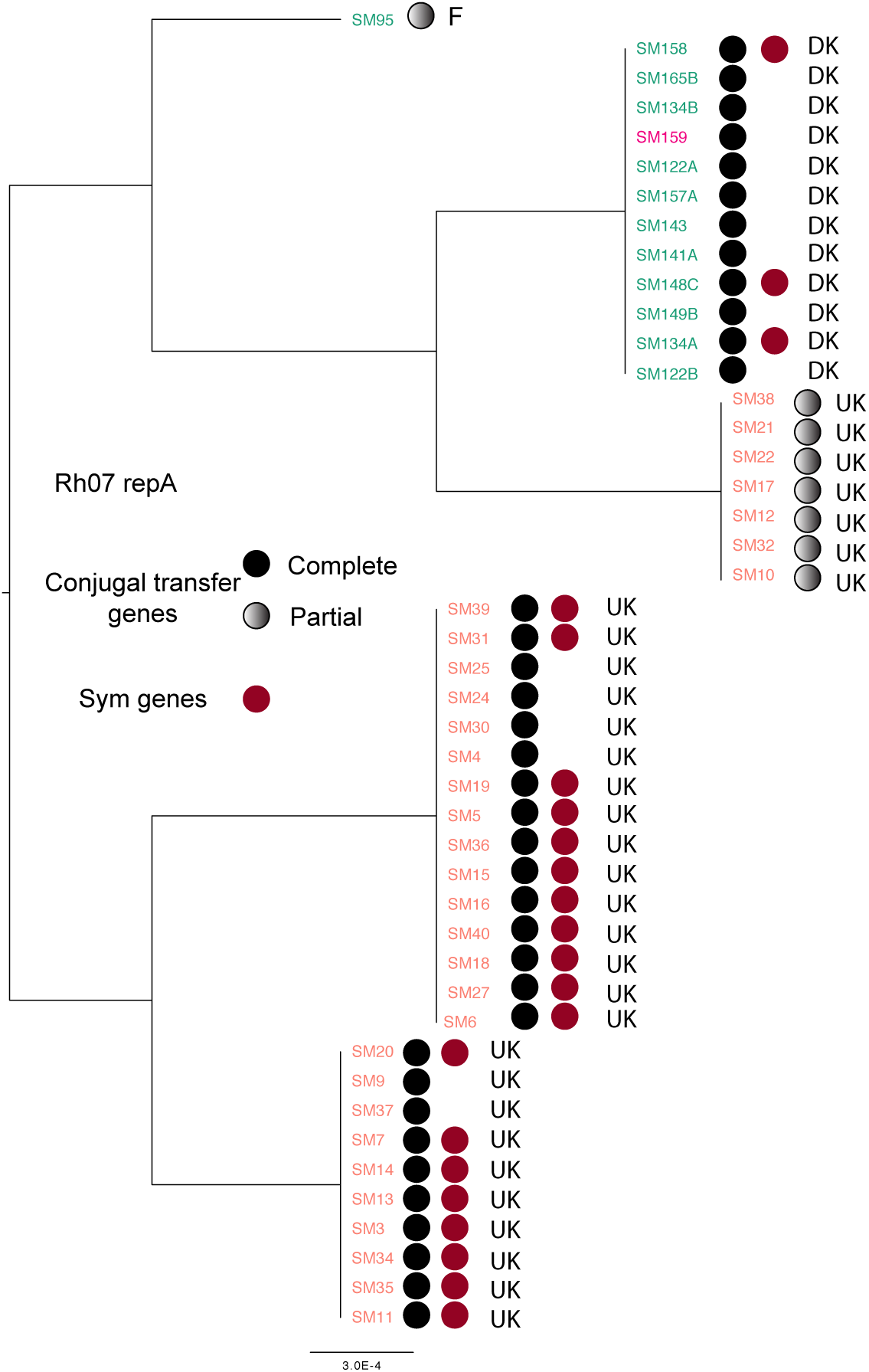
Phylogenetic analysis of the *repA* gene of plasmid type Rh07. DKO represents strains sampled from Danish organic fields, DK from Danish conventional trials. A complete set of conjugal transfer genes has the following genes upstream of *repA*: *traI*,*trbBCDEJKLFGHI*,*traRMHBFACDG*, with the origin of transfer (*oriT*) between *traA* and *traC*. Partial sets are broken by the end of the scaffold, mostly after *traM*.

**Figure S13:**
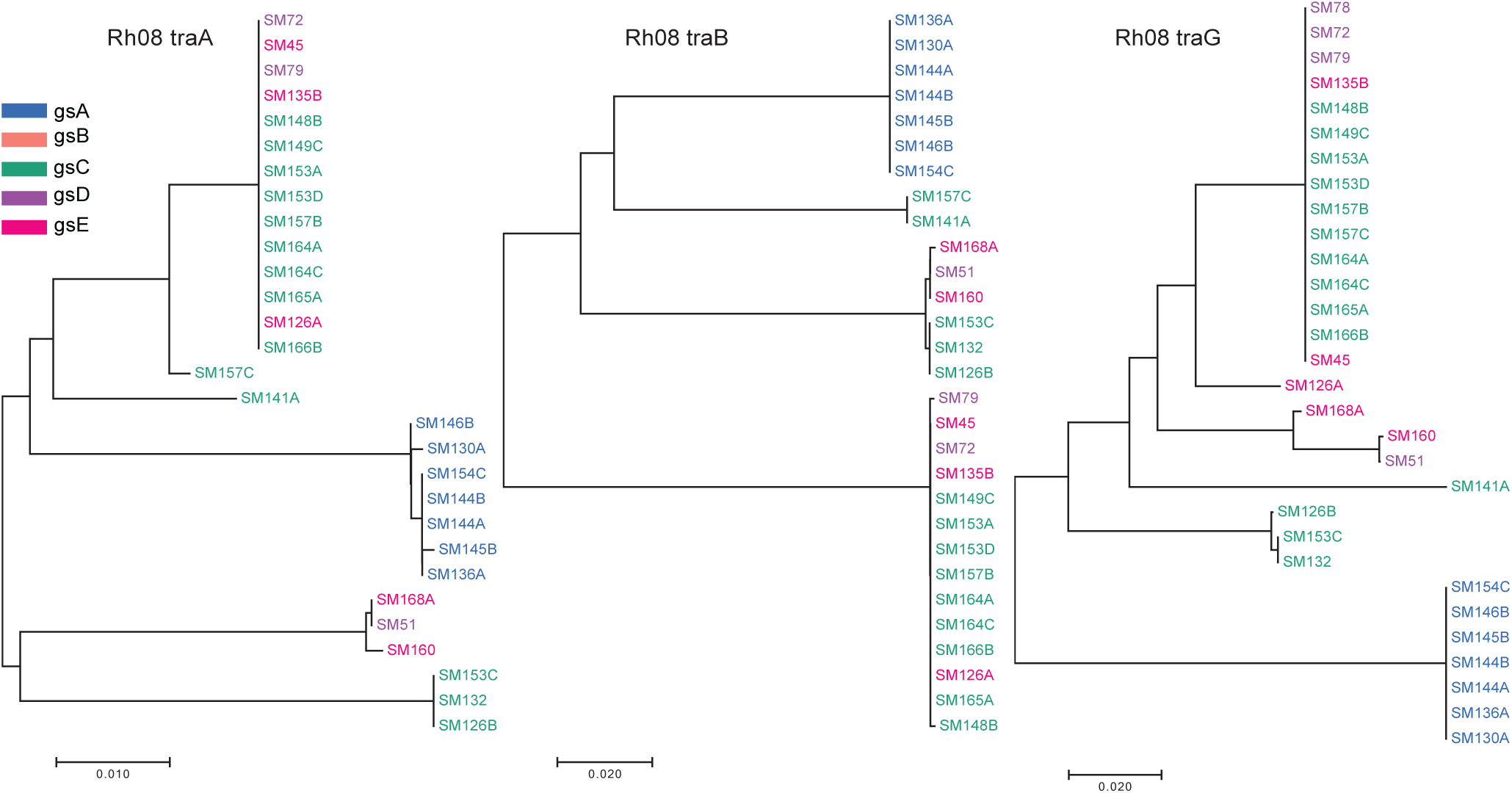
Phylogenetic trees of tranfer genes (*tra*) essential for the conjugation process. These genes are found in strains containing the plasmid Rh08, which is the sym-plasmid for some of the strains. A complete set of conjugal transfer genes has the following genes upstream of *repA*: *traI*,*trbBCDEJKLFGHI*,*traRMHBFACDG*.)

**Figure S14:**
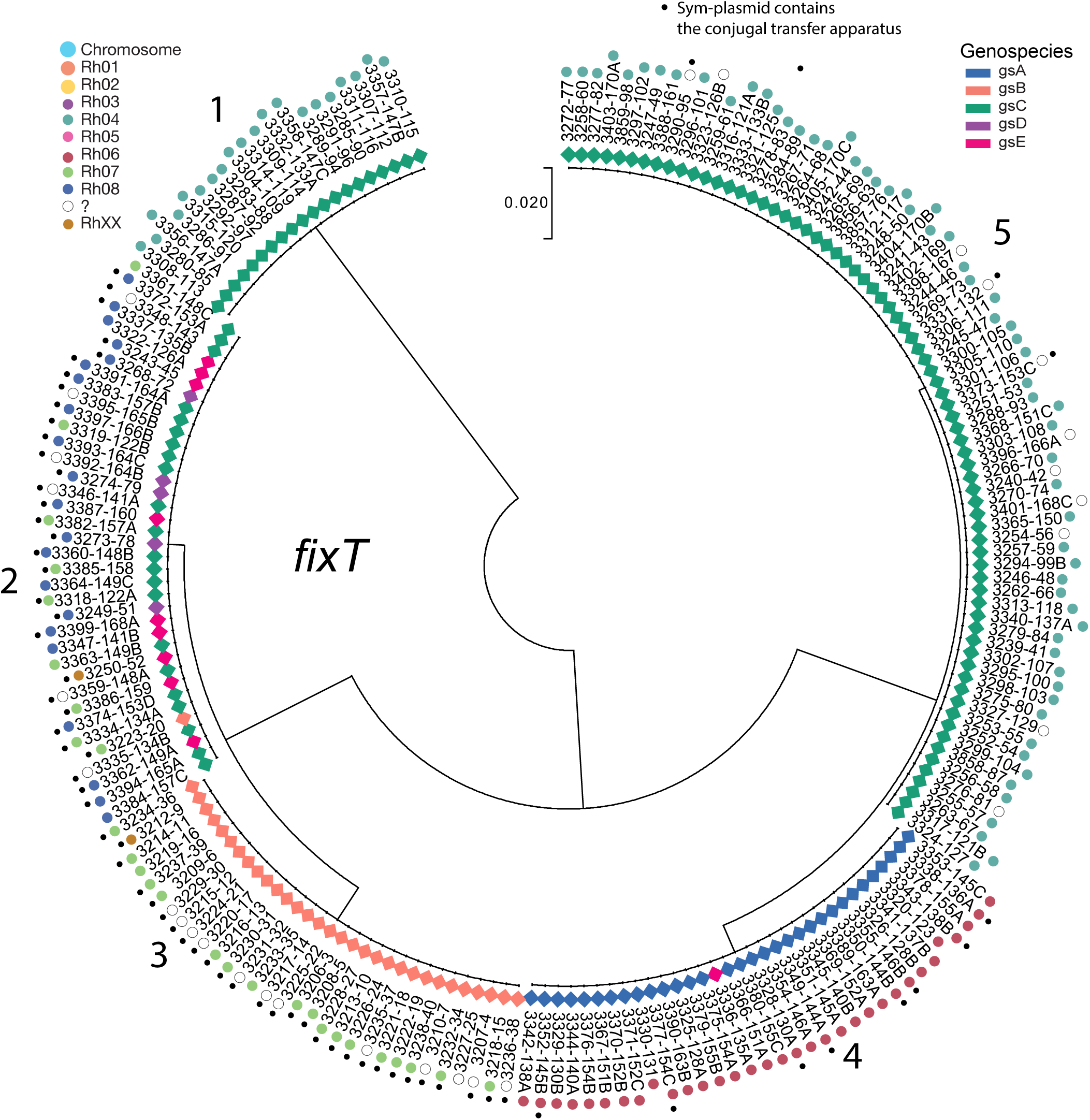
Phylogenetic tree of *fixT* and sym-plasmid classification. Dots correspond to strains containing a mobile sym-plasmid, with conjugal transfer system. With the exception of gsB clade (all strains from UK), no other clade is confined to a specific country of origin. All the numbers following the dash corresponds to the SM strain name.

## References

Alt-Mörbe, J et al. “The conjugal transfer system of Agrobacterium tumefaciens octopine-type Ti plasmids is closely related to the transfer system of an IncP plasmid and distantly related to Ti plasmid virgenes.” In: Journal of bacteriology 178.14 (1996), pp. 4248–4257.

Amarger, Nöelle and Lobreau, Jean Pierre. “Quantita-tive study of nodulation competitiveness in Rhizobium strains”. In: Appl. Environ. Microbiol. 44(3) (1982), pp. 583–588.

Andrews, Mitchell, et al. “Horizontal transfer of symbiosis genes within and between rhizobial genera: occurrence and importance”. In: Genes 9.7 (2018), p. 321.

Azad, Rajeev K and Lawrence, Jeffrey G. “Detecting laterally transferred genes: use of entropic clustering methods and genome position”. In: Nucleic acids research 35.14 (2007), pp. 4629–4639.

Bailly, Xavier, et al. “Population genomics of Sinorhizobium medicae based on low-coverage sequencing of sympatric isolates”. In: The ISME journal 5.11 (2011), p. 1722.

Bailly, Xavier, et al. “Recombination and selection shape the molecular diversity pattern of nitrogen-fixing Sinorhizobium sp. associated to Medicago”. In: Molecular Ecology 15.10 (2006), pp. 2719–2734.

Bankevich, Anton, et al. “SPAdes: a new genome assembly algorithm and its applications to single-cell sequencing”. In: Journal of computational biology 19.5 (2012), pp. 455–477.

Barnett, Melanie J, et al. “Nucleotide sequence and predicted functions of the entire Sinorhizobium meliloti pSymA megaplasmid”. In: Proceedings of the National Academy of Sciences 98.17 (2001), pp. 9883–9888.

Bever, JD. “Dynamics within mutualism and the maintenance of diversity: inference from a model of interguild frequency dependence”. In: Ecology Letters 2.1 (1999), pp. 52–61.

Burge, Christopher B and Karlin, Samuel. “Finding the genes in genomic DNA”. In: Current opinion in structural biology 8.3 (1998), pp. 346–354.

Cervantes, Laura, et al. “The conjugative plasmid of a bean-nodulating Sinorhizobium fredii strain is assembled from sequences of two Rhizobium plasmids and the chromosome of a Sinorhizobium strain”. In: BMC microbiology 11.1 (2011), p. 149.

Chen, Lishan, et al. “A new type IV secretion system promotes conjugal transfer in Agrobacterium tumefaciens”. In: Journal of bacteriology 184.17 (2002), pp. 4838–4845.

Daubin, Vincent, Lerat, Emmanuelle, and Perrière, Guy. “The source of laterally transferred genes in bacterial genomes”. In: Genome biology 4.9 (2003), R57.

Doolittle, W Ford. “Lateral genomics”. In: Trends in Biochemical Sciences 24.12 (1999), pp. M5–M8.

Friesen, Maren L. “Widespread fitness alignment in the legume–rhizobium symbiosis”. In: New Phytologist 194.4 (2012), pp. 1096–1111.

Galibert, Francis, et al. “The composite genome of the legume symbiont Sinorhizobium meliloti”. In: Science 293.5530 (2001), pp. 668–672.

Garud, Nandita R, et al. “Recent selective sweeps in North American Drosophila melanogaster show signatures of soft sweeps”. In: PLoS genetics 11.2 (2015), e1005004.

Greenlon, Alex, et al. “Global-level population genomics reveals differential effects of geography and phylogeny on horizontal gene transfer in soil bacteria”. In: Proceedings of the National Academy of Sciences 116.30 (2019), pp. 15200–15209.

Guillot, Gilles and Rousset, François. “Dismantling the Mantel tests”. In: Methods in Ecology and Evolution 4.4 (2013), pp. 336–344.

Gurevich, Alexey, et al. “QUAST: quality assessment tool for genome assemblies”. In: Bioinformatics 29.8 (2013), pp. 1072–1075.

Hanage, William P. “Not so simple after all: bacteria, their population genetics, and recombination”. In: Cold Spring Harbor perspectives in biology 8.7 (2016), a018069.

Harmon, Luke J and Glor, Richard E. “Poor statistical performance of the Mantel test in phylogenetic comparative analyses”. In: Evolution: International Journal of Organic Evolution 64.7 (2010), pp. 2173–2178.

Harrison, Peter W, et al. “Introducing the bacterial chromid: not a chromosome, not a plasmid”. In: Trends in microbiology 18.4 (2010), pp. 141–148.

Haukka, Kaisa, Lindström, Kristina, and Young, J Peter W. “Three phylogenetic groups of nodA and nifHGenes in Sinorhizobium and Mesorhizobium isolates from leguminous trees growing in Africa and Latin America”. In: Appl. Environ. Microbiol. 64.2 (1998), pp. 419–426.

Heidelberg, John F, et al. “DNA sequence of both chromosomes of the cholera pathogen Vibrio cholerae”. In: Nature 406.6795 (2000), p. 477.

Hirsch, PR, et al. “Physical Identification of Bacteriocinogenic, Nodulation and Other Plasmids in Strains of Rhizobium leguminosarum”. In: Microbiology 120.2 (1980), pp. 403–412.

Jain, Chirag, et al. “High throughput ANI analysis of 90K prokaryotic genomes reveals clear species boundaries”. In: Nature communications 9.1 (2018), p. 5114.

Konstantinidis, Konstantinos T, Ramette, Alban, and Tiedje, James M. “The bacterial species definition in the genomic era”. In: Philosophical Transactions of the Royal Society B: Biological Sciences 361.1475 (2006), pp. 1929–1940.

Konstantinidis, Konstantinos T and Tiedje, James M. “Prokaryotic taxonomy and phylogeny in the genomic era: advancements and challenges ahead”. In: Current opinion in microbiology 10.5 (2007), pp. 504–509.

Kumar, Nitin, et al. “Bacterial genospecies that are not ecologically coherent: population genomics of Rhizobium leguminosarum”. In: Open biology 5.1 (2015), p. 140133.

Laguerre, Gisèle et al. “Classification of rhizobia based on nodC and nifH gene analysis reveals a close phylogenetic relationship among Phaseolus vulgaris symbionts”. In: Microbiology 147.4 (2001), pp. 981–993.

Lawrence, Jeffrey G and Ochman, Howard. “Reconciling the many faces of lateral gene transfer”. In: TRENDS in Microbiology 10.1 (2002), pp. 1–4.

Lechner, Marcus, et al. “Orthology detection combining clustering and synteny for very large datasets”. In: PLoS One 9.8 (2014), e105015.

Lemaire, Benny, et al. “Symbiotic diversity, specificity 125 and distribution of rhizobia in native legumes of the Core Cape Subregion (South Africa)”. In: FEMS Microbiology Ecology 91.2 (2015), pp. 2–17.

Leplae, Raphaël et al. “Diversity of bacterial type II toxin–antitoxin systems: a comprehensive search and functional analysis of novel families”. In: Nucleic acids research 39.13 (2011), pp. 5513–5525.

Lerat, Emmanuelle, Daubin, Vincent, and Moran, Nancy A. “From gene trees to organismal phylogeny in prokaryotes: the case of the γ-Proteobacteria”. In: PLoS biology 1.1 (2003), e19.

Ling, Jun, et al. “Plant nodulation inducers enhance horizontal gene transfer of Azorhizobium caulinodans symbiosis island”. In: Proceedings of the National Academy of Sciences 113.48 (2016), pp. 13875–13880.

Long, Quan, et al. “Massive genomic variation and strong selection in Arabidopsis thaliana lines from Sweden”. In: Nature genetics 45.8 (2013), p. 884.

Mamidi, Sujan, et al. “Genome-wide association studies identifies seven major regions responsible for iron deficiency chlorosis in soybean (Glycine max)”. In: PloS one 9.9 (2014), e107469.

Mangin, Brigitte, et al. “Novel measures of linkage disequilibrium that correct the bias due to population structure and relatedness”. In: Heredity 108.3 (2012), p. 285.

Masuda, Hisako and Inouye, Masayori. “Toxins of prokaryotic toxin-antitoxin systems with sequence-specific endoribonuclease activity”. In: Toxins 9.4 (2017), p. 140.

Merkl, Rainer. “SIGI: score-based identification of genomic islands”. In: BMC bioinformatics 5.1 (2004), p. 22.

Nandasena, Kemanthi G, et al. “Rapid in situ evolution of nodulating strains for Biserrula pelecinus L. through lateral transfer of a symbiosis island from the original mesorhizobial inoculant”. In: Appl. Environ. Microbiol. 72.11 (2006), pp. 7365–7367.

Needleman, Saul B and Wunsch, Christian D. “A general method applicable to the search for similarities in the amino acid sequence of two proteins”. In: Journal of molecular biology 48.3 (1970), pp. 443–453.

Ochman, Howard, Lawrence, Jeffrey G, and Groisman, Eduardo A. “Lateral gene transfer and the nature of bacterial innovation”. In: nature 405.6784 (2000), p. 299.

Passel, Mark WJ van et al. “An acquisition account of genomic islands based on genome signature comparisons”. In: BMC genomics 6.1 (2005), p. 163.

Perez Carrascal, Olga M, et al. “Population genomics of the symbiotic plasmids of sympatric nitrogen-fixing Rhizobium species associated with Phaseolus vulgaris”. In: Environmental microbiology 18.8 (2016), pp. 2660–2676.

Provorov, NA, Andronov, EE, and Onishchuk, OP. “Forms of natural selection controlling the genomic 180 evolution in nodule bacteria”. In: Russian Journal of Genetics 53.4 (2017), pp. 411–419.

Provorov, Nikolai A and Vorobyov, Nikolai I. “Interplay of Darwinian and frequency-dependent selection in the host-associated microbial populations”. In: Theoretical population biology 70.3 (2006), pp. 262–272.

Provorov, Nikolai A and Vorobyov, Nikolai I. “Population genetics of rhizobia: construction and analysis of an infection and release model”. In: Journal of theoretical biology 205.1 (2000), pp. 105–119.

Remigi, Philippe, et al. “Symbiosis within symbiosis: evolving nitrogen-fixing legume symbionts”. In: Trends in microbiology 24.1 (2016), pp. 63–75.

Rogel, Marco A, Ormeno-Orrillo, Ernesto, and Romero, Esperanza Martinez. “Symbiovars in rhizobia reflect 195 bacterial adaptation to legumes”. In: Systematic and Applied Microbiology 34.2 (2011), pp. 96–104.

Rosini, Roberto, et al. “Identification of novel genomic islands coding for antigenic pilus-like structures in Streptococcus agalactiae”. In: Molecular microbiology 61.1 (2006), pp. 126–141.

Rousset, Francois. “Partial Mantel tests: reply to Castellano and Balletto”. In: Evolution 56.9 (2002), pp. 1874–1875.

Sauvage, Christopher, et al. “Genome-wide association 205 in tomato reveals 44 candidate loci for fruit metabolic traits”. In: Plant physiology 165.3 (2014), pp. 1120–1132.

Schulein, Ralf and Dehio, Christoph. “The VirB/VirD4 type IV secretion system of Bartonella is essential for establishing intraerythrocytic infection”. In: Molecular microbiology 46.4 (2002), pp. 1053–1067.

Seemann, Torsten. “Prokka: rapid prokaryotic genome annotation”. In: Bioinformatics 30.14 (2014), pp. 2068–2069.

Segovia, Lorenzo, Young, J Peter W, and Martínez-Romero, Esperanza. “Reclassification of American Rhizobium leguminosarum biovar phaseoli type I strains as Rhizobium etli sp. nov.” In: International Journal of Systematic and Evolutionary Microbiology 43.2 (1993), pp. 374–377.

Sievers, Fabian, et al. “Fast, scalable generation of high-quality protein multiple sequence alignments using Clustal Omega”. In: Molecular systems biology 7.1 (2011).

Silva, Claudia, et al. “Rhizobium etli and Rhizobium gallicum nodulate common bean (Phaseolus vulgaris) in a traditionally managed milpa plot in Mexico: population genetics and biogeographic implications”. In: Appl. Environ. Microbiol. 69.2 (2003), pp. 884–893.

Simonsen, Martin and Pedersen, Christian NS. “Rapid computation of distance estimators from nucleotide and amino acid alignments”. In: Proceedings of the 2011 ACM Symposium on Applied Computing. ACM. 2011, pp. 89–93.

Sukumaran, Jeet and Holder, Mark T. “DendroPy: a Python library for phylogenetic computing”. In: Bioinformatics 26.12 (2010), pp. 1569–1571.

Sullivan, John T, et al. “Comparative sequence analysis of the symbiosis island of Mesorhizobium loti 240 strain R7A”. In: Journal of Bacteriology 184.11 (2002), pp. 3086–3095.

Sullivan, John T, et al. “Nodulating strains of Rhizobium loti arise through chromosomal symbiotic gene transfer in the environment”. In: Proceedings of the National Academy of Sciences 92.19 (1995), pp. 8985–8989.

Tarjan, Robert. “Depth-first search and linear graph algorithms”. In: SIAM journal on computing 1.2 (1972), pp. 146–160.

Van Cauwenberghe, Jannick, et al. “Population structure of root nodulating Rhizobium leguminosarum in Vicia cracca populations at local to regional geographic scales”. In: Systematic and applied microbiology 37.8 (2014), pp. 613–621.

VanRaden, Paul M. “Efficient methods to compute genomic predictions”. In: Journal of dairy science 91.11 (2008), pp. 4414–4423.

Vuong, Holly B, Thrall, Peter H, and Barrett, Luke G. “Host species and environmental variation can influence rhizobial community composition”. In: Journal of Ecology 105.2 (2017), pp. 540–548.

Wetzel, Margaret E, et al. “The repABC plasmids with quorum-regulated transfer systems in members of the Rhizobiales divide into two structurally and separately evolving groups”. In: Genome biology and evolution 7.12 (2015), pp. 3337–3357.

Yoder, Jeremy B, et al. “Genomic signature of adaptation to climate in Medicago truncatula”. In: Genetics 196.4 (2014), pp. 1263–1275.

Young, J Peter W et al. “The genome of Rhizobium 270 leguminosarum has recognizable core and accessory components”. In: Genome biology 7.4 (2006), R34.

Zhao, Ran, et al. “Adaptive evolution of rhizobial symbiotic compatibility mediated by co-evolved insertion sequences”. In: The ISME journal 12.1 (2018), p. 101.

